# The HSF-DREB-MYB transcriptional regulatory module regulates flavonol biosynthesis and flavonoid B-ring hydroxylation in banana (*Musa acuminata*)

**DOI:** 10.1101/2023.08.23.554507

**Authors:** Jogindra Naik, Ruchika Rajput, Samar Singh, Ralf Stracke, Ashutosh Pandey

## Abstract

Plant flavonols act primarily as ultraviolet radiation absorbers, reactive oxygen species scavengers, and phytoalexins, and they contribute to biotic and abiotic stress tolerance in plants. Banana (*Musa acuminata*), an herbaceous monocot and important fruit crop, accumulates flavonol derivatives in different organs, including the edible fruit pulp. Although flavonol content varies greatly in different organs, the molecular mechanisms involving transcriptional regulation of flavonol synthesis in banana are not known. Here, we characterized three SG7-R2R3 MYB transcription factors (MaMYBFA1, MaMYBFA2, and MaMYBFA3) and their upstream regulators, heat shock transcription factor (MaHSF11) and dehydration responsive element binding factor (MaDREB1), to elucidate the molecular mechanism involved in transcriptional regulation of flavonol biosynthesis in banana. MaMYBFA positively regulate *flavonol synthase* 2 (*MaFLS2)* and downregulates *MaFLS1*. We show these transcription factors to be weak regulators of flavonol synthesis. Overexpression of *MaHSF11* enhances flavonol contents, particularly that of myricetin, and promotes flavonol B-ring hydroxylation, which contributes to the diversity of flavonol derivatives. MaHSF11 directly interacts with the *MaFLS1* and *flavonoid 3*′*, 5*′*-hydroxylase*1 (*MaF3*′*5*′*H1)* promoters, both *in vitro* and *in vivo*. MaHSF11 activates the expression of *MaDREB1* directly, which in turn regulates the expression of *MaMYBFA3*. Overall, our study elucidates a novel regulatory mechanism for flavonol synthesis in banana and suggests possible targets for genetic optimization to enhance nutritional value and stress responses in this globally important fruit crop.

**Significance statement:** Our study reveals that the three stress-responsive SG7 MYB factors, MaMYBFA1-3, are modest regulators of flavonol biosynthesis in banana. In contrast, the drought– and heat-responsive MaHSF11 plays a strong, positive role in this pathway, enhancing flavonol biosynthesis and flavonoid B-ring hydroxylation by upregulating key biosynthetic genes, such as *MaFLS1* and *MaF3’5’H1*, as well as *MaDREB1*, another regulator of flavonol biosynthesis in banana.

## Introduction

Plants produce a multitude of chemical compounds in response and adaptation to external and environmental stimuli, many of which are specialized metabolites (Oh *et al*., 2009; Mikkelsen *et al*., 2015; Murcia *et al*., 2017). Among these, flavonoids are ubiquitous and play vital roles in both regulation of developmental processes and defense against pathogens (Halbwirth *et al*., 2010; Maloney *et al*., 2014; Chen *et al*., 2019; Yang *et al*., 2021). Based on their structures, flavonoids are classified into flavonols, flavones, flavanones, isoflavones, anthocyanins, and proanthocyanidins (Lago *et al*., 2014). Flavonols have a basic 16-carbon, triple-ring structure containing a 3-hydroxyflavone backbone, and the different modifications and substituents in this ring give rise to their diversity (Ko *et al*., 2014). Besides acting as co-pigments with anthocyanins in flower petals and as reactive oxygen species (ROS) scavengers, flavonols have numerous other important functions in plants (Kitamura *et al*., 2016; Watkins *et al*., 2017). They absorb ultraviolet radiation (UV), protecting plants from UV-induced oxidative damage (Liang *et al*., 2020) and also regulate root growth and lateral root development by inhibiting polar auxin transport via stabilization of PIN-FORMED (PIN) protein dimers (Tan *et al*., 2019; Chapman *et al*., 2021; Teale *et al*., 2021). In addition, flavonols regulate pollen development and germination, nodule development, and symbiosis (Zhang *et al*., 2009; Muhlemann *et al*., 2018). Flavonols are synthesized by the flavonoid biosynthesis pathway involving the enzymes chalcone synthase (CHS), chalcone isomerase (CHI), flavanone 3-hydroxylase (F3H), flavonoid 3′ hydroxylase (F3′H), flavonoid 3′5′ hydroxylase (F3′5′H), and flavonol synthase (FLS), which are well characterized in many plant species (Nagamatsu *et al*., 2007; Guo *et al*., 2019; Jiang *et al*., 2020).

The flavonol biosynthesis pathway is regulated at several levels, and its transcriptional regulation has been extensively studied (Matus *et al*., 2009). A group of MYB transcription factors (TFs), known as subgroup 7 (SG7) R2R3-MYBs, play a major role in the regulation of flavonol biosynthesis (Schwinn *et al*., 2016); R2 and R3 are conserved regions. MYB12, MYB111, and MYB11, also known as PRODUCTION OF FLAVONOL GLYCOSIDES1–3 (PFG1–3), respectively, modulate flavonol biosynthesis in a tissue-specific manner in Arabidopsis (*Arabidopsis thaliana*) (Mehrtens *et al*., 2005; Stracke *et al*., 2007; Stracke *et al*., 2010), tobacco (*Nicotiana tabacum*) (Misra *et al*., 2010; Pandey *et al*., 2014a, b; Pandey *et al*., 2015a), and tomato (*Solanum lycopersicum*) (Pandey *et al*., 2015b). They regulate the expression of several rate-limiting biosynthesis genes, including *CHS*, *F3H*, and *FLS*. *FLS* is also coordinately regulated by MYB99, MYB21, and MYB24 in Arabidopsis stamens and anthers independently of the PFGs (Battat *et al*., 2019; Zhang *et al*., 2021). Many MYB TFs belonging to SG7 have been reported to regulate flavonol synthesis in various species namely VvMYBF1 (*Vitis vinifera*), CmMYB3 (*Chrysanthemum morifolium*), MtMYB134 (*Medicago truncatula*), CaMYB39 (*Cicer arietinum*), MdMYB22 (*Malus domestica*), NtMYB12 (*Nicotiana tabacum*) are among the several factors characterized so far to regulate the expression of *FLS* and flavonol synthesis (Czemmel *et al*., 2009; Wang *et al*., 2017; Song *et al*., 2020; Naik *et al*., 2021; Saxena *et al*., 2023; Yang *et al*., 2023). Apart from these, AtHY5, AtARF2, AtWRKY23, NtWRKY11b, VvWRKY70, GmNAC2, MaABI5-like, MaC2H2, PaNAC03, CsNAC086 are also among the few transcriptional regulators of flavonol synthesis in different species including banana (Grunewald *et al*., 2012; Dalman *et al*., 2017; Bian *et al*., 2020; Song *et al*., 2020; Bhatia *et al*., 2021; Jiang *et al*., 2022; Song *et al*., 2022; Song *et al*., 2023; Wang *et al*., 2021f; Wei *et al*., 2023b; Song *et al*., 2024). A recent study highlighted rather an interesting mechanism, where a SG7 MYB ScMYB12 interacts with ScTT8 to regulate flavonol synthesis in Jojoba (Zheng *et al*., 2024). DELLA component (RGA and GAI) directly interacts with MYB12 and MYB111 to promote flavonol synthesis in Arabidopsis (Tan *et al*., 2019). Many of these characterized factors regulate flavonol biosynthesis pathway in hormone– and stress-responsive manner.

Because plants are exposed to numerous biotic and abiotic stresses, such as excessive light and UV radiation, suboptimal water conditions (drought and waterlogging), high salt, extreme temperatures, nutrient deficiencies, and pathogen infection (López *et al*., 2008; Cramer *et al*., 2011; Kalaji *et al*., 2017; Choudhury *et al*., 2017), they have evolved various defensive mechanisms that are activated after exposure to stress and involve a complex interplay of signaling pathways. Other classes of TFs, such as heat shock factors (HSFs) and dehydration-responsive element-binding (DREB) factors, modulate the plant’s stress response to drought and high heat (Zhao *et al*., 2010; Lee *et al*., 2010; Huang *et al*., 2016; Zinsmeister *et al*., 2020). HSFs, in their dimeric form, regulate several stress-responsive genes by binding to heat shock elements (HSEs; nGAAnnTTCn/nTTCnnGAAn) via their N-terminal DNA-binding domains (Liu *et al*., 2008; Wang *et al*., 2020). HSFs can be classified into three major classes, A, B, and C, depending upon the type of amino acid residues and the length of flexible linker length between DBD (DNA-binding domain) and OD (oligomerization domain) (Scharf *et al*., 2012; Manna *et al*., 2021). In addition to heat stress tolerance, HSFs regulate salt, drought, and cold stress responses in Arabidopsis (Yoshida *et al*., 2011). MdHSFA8a is a positive regulator of drought stress tolerance and flavonoid synthesis in apple (*Malus domestica*) (Wang *et al*., 2020). GmHSFB2b positively regulate flavonol synthesis in soybean through up-regulation of *GmCHS* and down-regulation of *GmNAC2*, a repressor of *GmFLS1* and *GmFLS2* under salt stress (Bian *et al*., 2020). Apart from these, The HSFA2 and HSFA3 heterodimeric complex confers transcriptional thermal memory by promoting epigenetic H3K4 hypermethylation (Friedrich *et al*., 2021). In banana, MaHSF24 negatively regulates cold stress response via repression of HSP and antioxidant genes, while MaHSF26 alleviates chilling injury in heat shock responsive manner via repressing MaRBOH (Si *et al*., 2022; Si *et al*., 2024). DREBs, which belong to the AP2/ERF family of TFs acting both upstream and downstream of HSF, are also involved in several stress signaling pathways, in particular drought, salt, heat, and cold stress (Dubouzet *et al*., 2003; Schramm *et al*., 2008; Chen *et al*., 2016; Huang *et al*., 2016). CBF/DREB TFs bind to the cold and dehydration-responsive CRT/DREB motif core element (CCGAC) present in the promoters of their target genes via the AP2/ERF DNA-binding domain (Li *et al*., 2020; Wang *et al*., 2021a). In context of metabolic pathways regulation, they are known to influence the expression of flavonoid related genes in different plants. For instance, in eggplant, SmCBF1/2/3 interacts with SmMYB113 and exerts additive effect on the SmMYB113 and SmTT8 mediated activation of anthocyanin synthesis (Zhou *et al*., 2020). CitERF32/33 and CitRAV1 cooperatively promote citrus flavone synthesis (Zhao *et al*., 2021). MdAP2-34, a DREB, modulates flavonol content in apple (Han *et al*., 2022). In addition, DREBs regulate fruit ripening via a complex transcriptional regulatory pathway in banana (*Musa acuminata*) (Kuang *et al*., 2017). MaCBF1 interacts with MaNAC1, a downstream target of MaICE1, to confer cold stress tolerance in banana (Shan *et al*., 2014). Among all the abiotic stress, high temperature is also an important factor where HSFs are master downstream components in heat stress response (Liu *et al*., 2012; Guo *et al*., 2016; Rao *et al*., 2021). It also poses a threat to the normal physiology of plants and greatly affects crop yield and production (Fahad *et al*., 2017). High temperature is known to modulate flavonoid accumulation in different plant species to confer heat tress tolerance. For instance, in oriental hybrid lily, LhMYBC2 negatively regulates anthocyanin biosynthesis in high temperature-responsive manner via competition with LhMYBA1 for LhTT8 (Yang *et al*., 2024). Likewise, an atypical SG7 R2R3 MYB CmMYB012 suppresses flavone and anthocyanin accumulation under high temperature in chrysanthemum (*Chrysanthemum x morifolium*) (Zhou *et al*., 2021). Whereas in *C. sinensis*, heat induced CsHSFA1b and CsHSFA2 activates CsJAZ6, which interacts with CsEGL3 and CsTTG1 to inhibit catechin synthesis under high temperature (Zhang *et al*., 2023). Theses finding indicates decreased accumulation of anthocyanin and proanthocyanidin under heat stress. On the contrary, high temperature promotes flavonol biosynthesis (Martinez *et al*., 2016; Fragkostefanakis *et al*., 2016; Paupière *et al*., 2017). Flavonols scavenge excessive ROS during heat stress to promote pollen viability and pollen tube growth in tomato (Muhlemann *et al*., 2020). In tomato, heat stress significantly enhanced accumulation of flavonols content in leaves and pollen (Martinez *et al*., 2016, Paupiere *et al*., 2017).

In terms of global production and monetary value, banana is the fourth most important food crop in the world, after rice (*Oryza sativa*), wheat (*Triticum aestivum*), and maize (*Zea mays*) (Wang *et al*., 2021c). Banana is a monocarpic herbaceous crop with climacteric fruits, which is characterized by postharvest ripening because of cellular respiration and increased ethylene production. However, global gross production is often affected by stresses including drought, cold, salinity, and, most importantly, *Fusarium oxysporum* f. sp. *cubense* 4 (Foc4), which causes a devastating wilt disease (Xu *et al*., 2010). We identified and functionally characterized three SG7 R2R3-MYB TFs (MaMYBFA1, MaMYBFA2, and MaMYBFA3) as regulators of flavonol synthesis in banana. These MaMYBFAs positively regulate *MaFLS2* and downregulate *MaFLS1*. MaHSF11 directly regulates the expression of *MaFLS1*, *MaF3*′*5*′*H1*, and *MaDREB1*. MaDREB1, in turn, regulates the expression of *MaMYBFA3*. Additionally, we identified MaHSF11 as a regulator of cytochrome P_450_ (CYP450) B-ring hydroxylase in banana, where it binds directly to the promoter of *MaF3*′*5*′*H1.* Taken together, our results elucidate a novel regulatory cascade of flavonol synthesis in the important fruit crop banana.

## Results

### An MaMYBFA1–3 triad modulates flavonol biosynthesis in banana

Our previous *in silico* study (Pucker *et al*., 2020) determined that the banana genome contains 285 *R2R3-MYB* genes. Three of the encoded MaMYBs form a clade with previously characterized flavonol regulators in a phylogenetic tree (Supplemental Fig. S1a). We selected these three candidates for further study and named them *MaMYBFA1* (Ma02_g00290), *MaMYBFA2* (Ma05_g23640), and *MaMYBFA3* (Ma08_g10260), where the abbreviation FA stands for flavonol activator. A multiple protein sequence alignment identified the conserved R2R3-MYB domain and the two SG7-specific, conserved motifs: SG7-1 and SG7-2 (Supplemental Fig. S1b). The full-length coding sequences (CDSs) of *MaMYBFA1*, *MaMYBFA2*, and *MaMYBFA3* were PCR-amplified from banana cDNA and used to express the encoded proteins. The subcellular localization of MaMYBFA-YFP fusion proteins was analyzed in *Nicotiana benthamiana* epidermal cells and was restricted to the nucleus, co-localizing with the nuclear marker protein NLS-RFP (Supplemental Fig. S1c), indicating that MaMYBFA1–3 are nuclear proteins.

We used RT-qPCR to analyze the expression of these three *MaMYBFA* genes in different organs/tissues of banana variety ‘Grand Naine’ (Supplemental Fig. S2a). All three *MaMYBFA* genes were differentially expressed in the analyzed organs. *MaMYBFA1* was most highly expressed in young leaf followed by bract tissue, and ripe peel. *MaMYBFA2* transcript levels were highest in young leaf followed by bract and ripe peel, and *MaMYBFA3* was expressed most highly in young leaf followed by bract and pseudostem. Transcript accumulation of the three genes was lowest in ripe pulp. These expression patterns suggest functional differences between the three MaMYBFAs in terms of upstream regulation and downstream regulatory potential toward biosynthesis genes. We also analyzed the expression level of *MaFLS1* and *MaFLS2*, the two probable targets of MaMYBFAs. The expression level of *MaFLS1* was highest in bract and young leaf, while the pseudostem, unripe and ripe peel, and unripe pulp had lower expression level (Supplemental Fig. S2b). The root and ripe pulp had least expressions among all the tissues. On the other hand, the expression level of *MaFLS2* was considerably higher in bract tissue only (Supplemental Fig. S2b). All other tissues, however, had very lower transcript level of *MaFLS2* with almost negligible expression in unripe peel (Supplemental Fig. S2b). Finally, we quantified different flavonol aglycones to study the differential tissue specific accumulation pattern. We found that kaempferol content was highest in leaf, unripe peel, and pulp. Likewise, quercetin content was highest in bract followed by other tissues, while myricetin content was higher in peel, bract and pseudostem tissue (Supplemental Fig. S2c). Banana ripe pulp tissue, however, accumulate very least amount of all the flavonol aglycones. This varying accumulation of *MYB* genes, *FLS* genes, and flavonol metabolites suggests multilayer regulatory mechanism operating to impact overall flavonol accumulation in banana.

To analyze the regulatory role of MaMYBFA1–3, we transiently overexpressed the *MaMYBFA1–3* CDSs under the control of the *Zea mays ubiquitin1* promoter (*proZmUBI1*) in immature banana fruit discs (Fig. 1a-b). The activity of a GUS reporter simultaneously transformed on the same T-DNA in immature banana fruit discs indicated successful transformation of the fruit tissues (Fig. 1b). RT-qPCR analysis revealed upregulation of the respective *MaMYBFA* transcripts in transgenic tissue samples (Fig. 1c). The expression of selected flavonoid biosynthesis genes was analyzed in MaMYBFA-overexpressing (OE) fruit discs. The expression of *CHALCONE SYNTHASE* (*MaCHS1–6*), *CHALCONE ISOMERASE* (*MaCHI1–2*), *FLAVANONE 3-HYDROXYLASE* (*MaF3H1–2*), and *FLAVONOL SYNTHASE* (*MaFLS1–2*) was modulated to different degrees in the OE fruit discs (Fig. 1d, Supplemental Figure S3). *MaMYBFA1* overexpression enhanced the expression of *MaCHS1*, whereas *MaMYBFA2* and *MaMYBFA3* overexpression did not (Supplemental Fig. S3). *MaCHS4* expression increased significantly with overexpression of all three *MaMYBFA*s. However, *MaCHS3* and *MaCHS5* showed enhanced expression only with overexpression of *MaMYBFA1* and *MaMYBFA3* (Supplemental Fig. S3), and only *MaMYBFA1* and *MaMYBFA2* overexpression increased the *MaCHS6* transcript level. *MaCHI* and *MaF3H* transcripts were also modulated in the transgenic OE fruit tissues: The highest activation of *MaCHI1* resulted from *MaMYBFA1* overexpression, whereas *MaCHI2* transcripts were increased 2-fold by *MaMYBFA3* overexpression (Supplemental Fig. S3). For *MaF3H1*, a significant increase in the transcript level was observed with *MaMYBFA1* and *MaMYBFA3* overexpression (Supplemental Fig. S3). Genes encoding CYP450 proteins also showed differential modulation; only *MaF3*′*5*′*H3* and *MaF3*′*5*′*H4* transcripts were upregulated in OE fruit tissues, whereas *MaF3*′*H* and other *MaF3*′*5*′*H* paralogs showed a marked decrease in expression (Supplemental Fig. S3). Finally, we analyzed the expression of *MaFLS1* and *MaFLS2*, the putative primary targets of MaMYBFAs, in transgenic *MaMYBFA* OE fruit discs (Fig. 1d). *MaFLS1* was significantly downregulated, whereas *MaFLS2* transcript levels increased in the *MaMYBFA* OE tissues (Fig. 1d). Overall, we found that overexpression of *MaMYBFA1* and *MaMYBFA3* significantly increased the expression of most structural flavonoid biosynthesis genes, whereas *MaMYBFA2* overexpression either decreased or negligibly affected the transcript levels of most of the genes studied.

**Figure 1.**
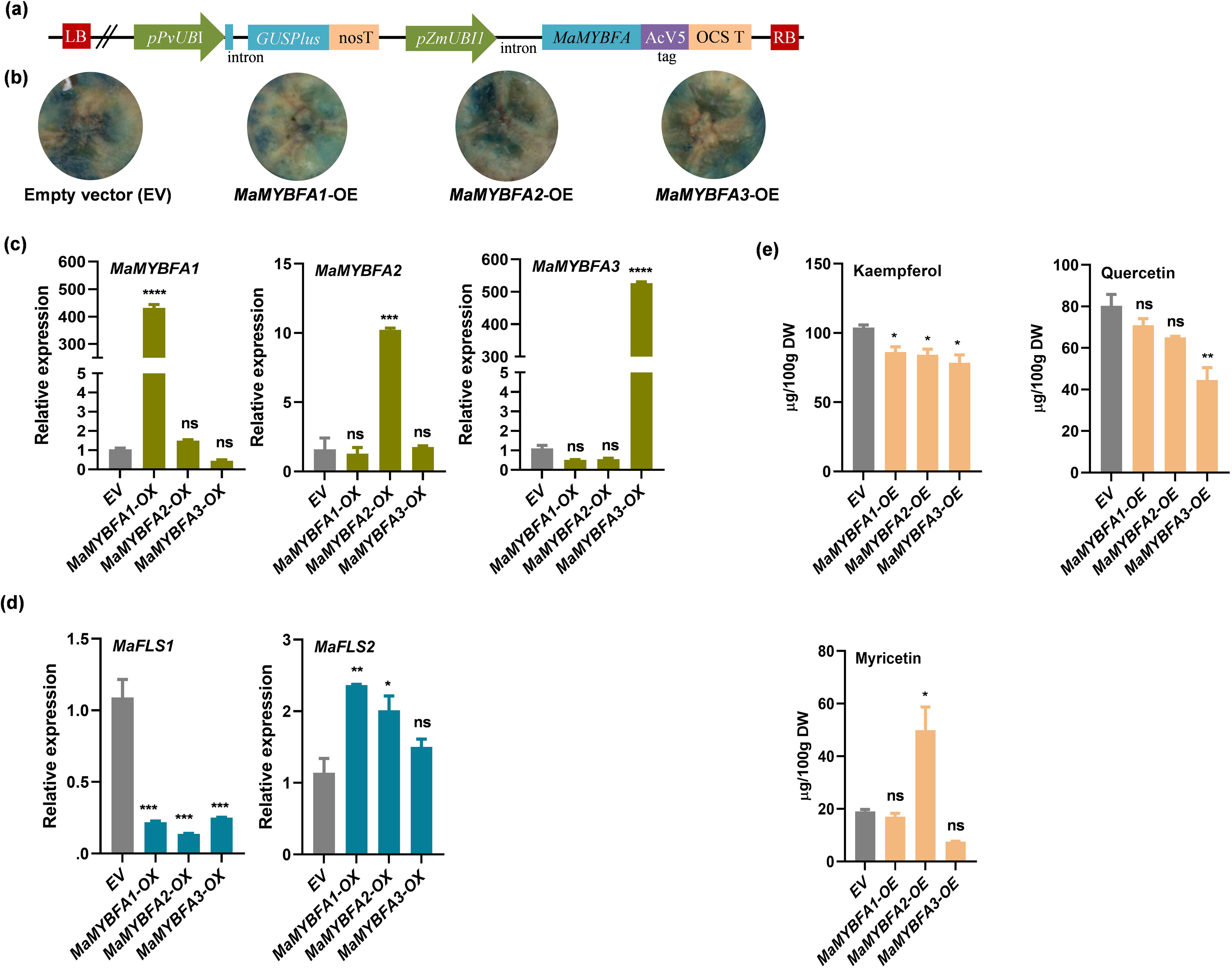
MaMYBFA1, MaMYBFA2, and MaMYBFA3 differentially regulate the expression of *MaFLS1* and *MaFLS2* and modulate flavonol biosynthesis in banana. (a) Diagram of the plant expression cassette used for transient overexpression (OE) of *MaMYBFA* in banana fruit discs. The *Zea mays ubiquitin1* promoter (*proZmUBI1*) drives the expression of *MaMYBFA*. The expression of the *GUSPlus* marker gene is driven by the *Phaseolus vulgaris ubiquitin1* promoter (*proPvUBI*). (b) Representative images showing GUS staining of transiently transformed banana fruit discs indicating successful transformation. Images were taken 3 days after transformation. (c-d) Relative expression of *MaMYBFA1–3* (c) and *MaFLS1* and *MaFLS2* (d) OE lines in transiently transformed *MaMYBFA* OE banana fruit disc tissues as determined by RT-qPCR. EV, pANIC6b empty vector control. Values are represented as fold change in three replicates of pool of 4-5 independent biological samples. (e) Flavonol quantification showing differential alteration in the accumulation of kaempferol, quercetin, and myricetin aglycones in the *MaMYBFA* OE banana fruit discs. Values are given in µg per 100 g of dry weight (DW) of fruit discs of three replicates. One-way ANOVA with Tukey’s multiple comparison was used for statistical analyses. *\*P*□≤□0.05; *\*\*P*□≤□0.01; *\*\*\*P*□≤□0.001; *\*\*\*\*P* ≤ 0.0001; ns, not significant.

To analyze the modulation of flavonol metabolite contents in transgenic *MaMYBFA* OE fruit disc tissues, we performed LC-MS-based, targeted flavonol quantification (Fig. 1e). We found decreased levels of kaempferol and quercetin in all the *MaMYBFA* OE tissues and an increase in myricetin content in *MaMYBFA2* OE compared to the empty vector (EV) control. Although the observed changes were significant for kaempferol, the contents of quercetin and myricetin were not altered significantly by *MaMYBFA1* and *MaMYBFA3* OE. Only *MaMYBFA2* OE significantly increased the accumulation of myricetin (Fig. 1e). Taken together, these results indicate that MaMYBFA1, MaMYBFA2, and MaMYBFA3 act as a MYB regulatory triad for modulating the expression of flavonol synthesis genes in banana. Despite significant up-regulation of *MaMYBFA* genes after their overexpression in banana fruits, flavonol content remain mostly unaltered or else there was a little change in the overall flavonol pool (Fig. 1e). This finding can be attributed to the downregulation of *MaFLS1*, *MaF3’H* and a few other genes, while increased expression of *MaFLS2* could compensate it.

### MaHSF11 promotes flavonol synthesis in banana by transactivating *MaFLS1* and regulating several other biosynthesis genes

None of the banana SG7 R2R3-MYBs we examined upregulated the expression of *MaFLS1* but instead repressed it. Because *MaFLS1* is highly expressed in leaves and is the major enzyme for flavonol synthesis in banana (Pandey *et al*., 2016; Busche *et al*., 2021), we attempted to identify a putative regulator of *MaFLS1*. An ∼2-kb promoter fragment was searched for possible TF binding sites, including ACGT-motif, E-box, LTRE (DRE/CRT), HSE, and W-box elements, which are recognized by bZIP, MYC, DREB/CBF, HSF, and WRKY factors, respectively (Supplemental File S1). Among these, the HSF TFs are poorly characterized in the context of flavonoids. The *MaFLS1* promoter contained two HSEs (nTTCnnGAAn) at positions −310 (HSE1) and −1154 (HSE2) upstream of the transcription start site (TSS). To identify a HSF as a possible HSE-binding candidate, we searched the publicly available banana transcriptome datasets related to abiotic stresses because both *HSF* and *FLS* are well-known stress-responsive genes. We found a transcriptome dataset of the drought-stressed banana cultivars Saba (ABB, drought tolerant) and Grand Naine (AAA, drought susceptible) to be the most suitable for this analysis (Muthusamy *et al*., 2016). From the transcriptome deep sequencing data, we found five *HSF* genes, *MaHSF9* (*Ma08_g28350*), *MaHSF11* (*Ma06_g33440*), *MaHSF10* (Ma03_g17030), *MaHSF25* (*Ma01_g03200*), and *MaHSF26* (Ma01_g09560), that were differentially expressed before and after drought treatment in the cultivar Grand Naine (Fig. 2a). *MaHSF25* and *MaHSF26* are class B HSFs, which are known to repress gene expression (Scharf *et al*., 2012). However, *MaHSF9*, *MaHSF10*, and *MaHSF11* are class A HSFs, which are known to act activate gene expression due to the AHA activator motif (Scharf *et al*., 2012).

**Figure 2.**
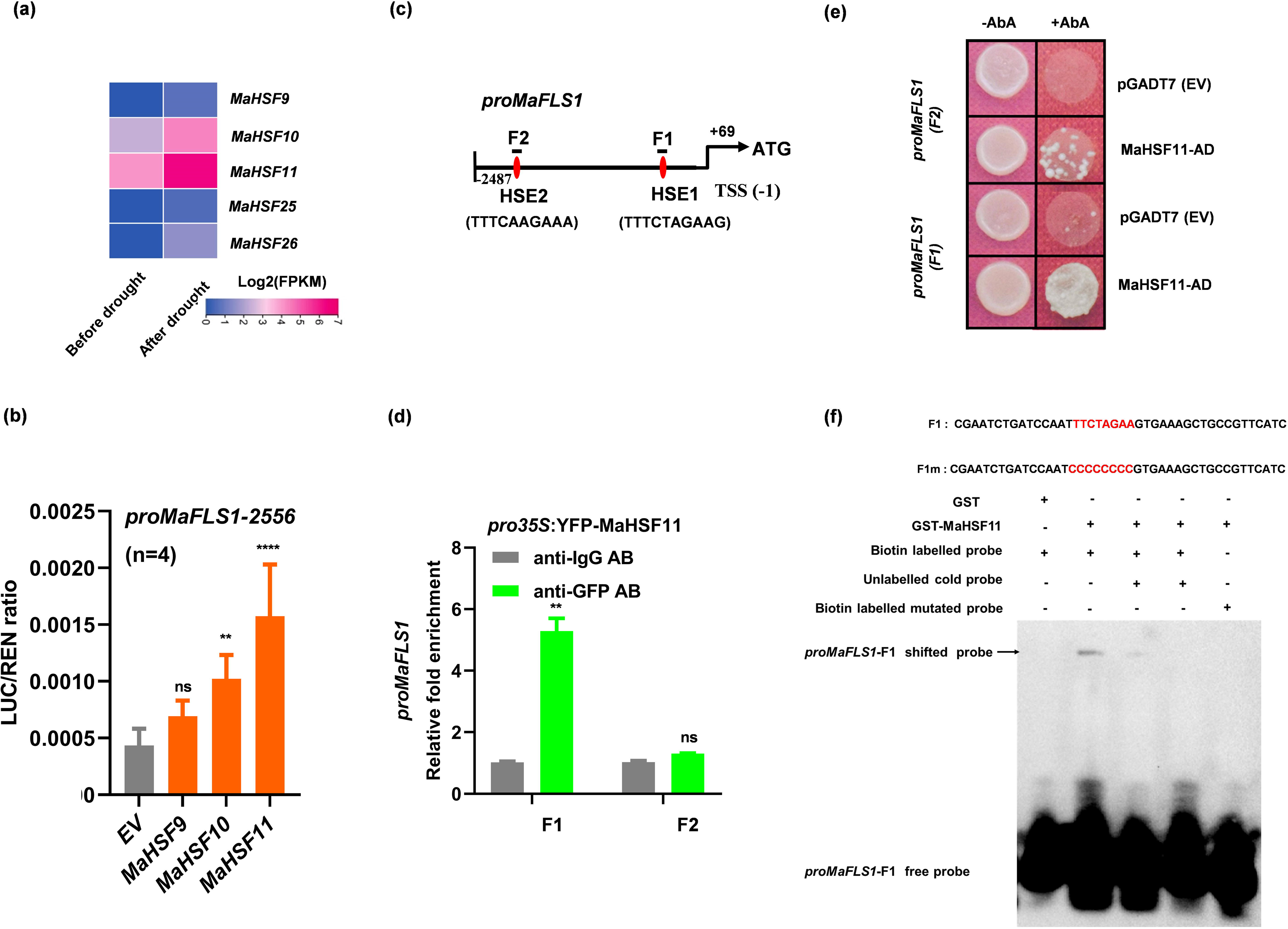
MaHSF11 directly binds to the *MaFLS1* promoter and regulates the expression of *MaFLS1* to modulate flavonol biosynthesis in banana. (a) Heatmap showing expression of five banana candidate *HSF* genes before and after drought treatment (Grand Naine cultivar, AAA genome). FPKM values for the indicated genes were extracted from publicly available RNA-Seq datasets and are represented according to the 0–7 color scale shown. (b) Dual-luciferase assay showing transactivation of *proMaFLS1–2556* by MaHSF. Values are shown as means ± SD of the LUC/REN ratios of four independent biological replicates (*n* = 4). (c) Illustration of the *cis*-motifs in the region located within 2487 bp upstream of the TSS in the *MaFLS1* promoter. F1 and F2 fragments contain one HSE element each having TTTCTAGAAG and TTTCAAGAAA motifs. +69 shows the bp distance between TSS and start codon. (d) ChIP-qPCR assay analyzing the binding of MaHSF11 to fragments F1 and F2 of *proMaFLS1*, both of which contain the HSE consensus sequence (nTTCnnGAAn). Fold enrichment values are given as means ± SD of two biological replicates. AB, antibody. (e) Y1H assay showing interaction of MaHSF11 with the two HSE-containing *proMaFLS1* fragments F1 and F2. The empty pGADT7 vector (EV) was used as a negative control. Interaction was indicated by yeast growth on synthetic Ura– dropout media supplemented with AbA. AD, activation domain. (f) EMSA showing the binding of MaHSF11-GST protein with the F1 fragment of *MaFLS1* promoter (*MaFLS1*-F1) *in vitro*. Binding cis-motif and the mutated motif in the probes are highlighted in red in F1 and F1m, respectively. MaHSF11-GST protein, labelled and unlabeled probes were separated in polyacrylamide gel, blotted on nylon membrane, and detected by chemiluminescence. + indicates presence and – indicates absence of probe/protein.

Using a dual-luciferase assay in *N. benthamiana* leaves, we examined whether MaHSF9, MaHSF10, and MaHSF11 could regulate the expression of *MaFLS1* by testing their ability to regulate *proMaFLS1–2556*. We observed a significant increase in the activity of *proMaFLS1–2556* when co-transfected with either *MaHSF10* or *MaHSF11* compared to the EV (Fig. 2b). Between these two, MaHSF11 showed the higher *in vivo* transactivation ability (>3.6-fold, *P* < 0.0001). Therefore, we focused on MaHSF11 in further studies and tested whether MaHSF11 could directly interact with *proMaFLS1*. To verify direct binding of MaHSF11 to the HSEs in the *MaFLS1* promoter, we performed a ChIP-qPCR assay in which *proMaFLS1* was co-infiltrated with a *pro35S:YFP-MaHSF11* construct into *N. benthamiana* leaves and expressed for 3 days. An anti-GFP antibody was used to immunoprecipitate YFP-MaHSF11 and the associated chromatin. A qPCR assay with purified DNA from the precipitated chromatin showed an approximate 5-fold enrichment of fragment F1 containing HSE1; fragment F2, containing HSE2, was not enriched (Fig. 2c,d). We then performed Y1H assays to validate the direct binding of MaHSF11 to HSEs of *proMaFLS1* using 100-bp, HSE-containing promoter fragments F1 and F2 to test for interaction with the MaHSF11-AD fusion protein. The Y1H growth assay showed strong binding of MaHSF11 to F1 and a weak interaction with F2 (Fig. 2e). Furthermore, we performed an EMSA experiment, which demonstrated the direct binding of MaHSF11-GST with the promoter fragment F1 of the *MaFLS1* promoter *in vitro*. MaHSF11 could bind to the TTCTAGAA motif-containing F1 fragment, but not with F1m, which has a mutated motif (Fig. 2c,f). Taken together, our results show that MaHSF11 binds directly to the HSEs in the promoter of *MaFLS1* and activates its expression.

Phylogenetic analysis revealed clustering of MaHSF11 with AtHSFA4a, which confers salt tolerance in the MPK3/6-dependent pathway in Arabidopsis, and other A-class HSF proteins (Fig. 3a). Furthermore, a subcellular localization assay in *N. benthamiana* leaves using a pro35S:YFP-MaHSF11 construct confirmed nuclear localization of YFP-MaHSF11, which co-localized with a nuclear NLS-RFP marker (Fig. 3b). In addition to nuclear localization, we also observed fluorescence in plasma membrane, suggesting probable post-translational modification (Fig. 3b). In tissue-specific expression pattern analysis, we found that *MaHSF11* is highly expressed in roots and leaves, followed by other vegetative tissues. Meanwhile, ripe peel and ripe pulp have very low transcript levels. (Fig. 3c).

**Figure 3.**
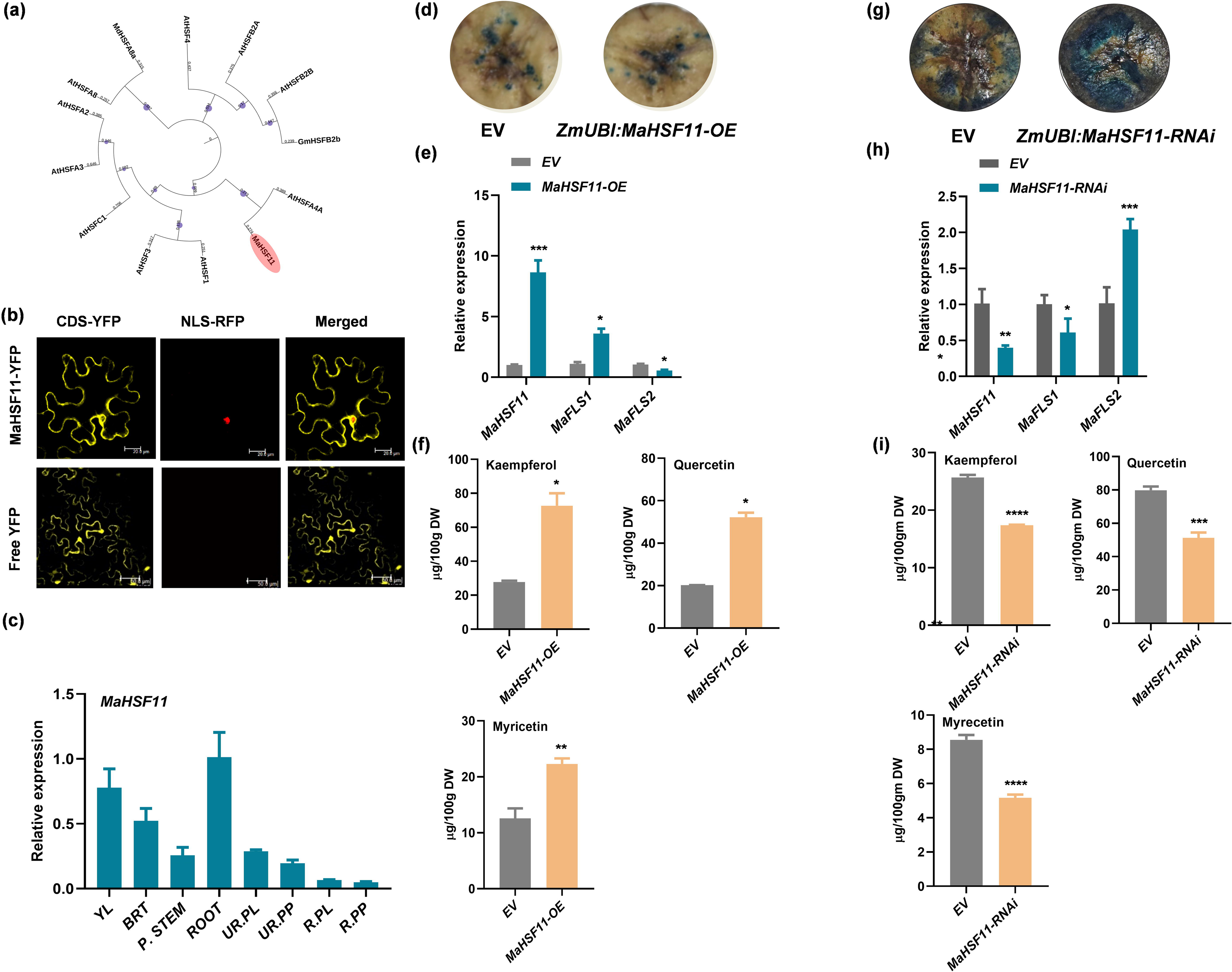
MaHSF11 promotes flavonol synthesis in banana fruit. (a) The phylogenetic tree was constructed in MEGA-X32 software using the maximum-likelihood method with 1000 bootstrap values and was visualized with iTOL v.6.4 (http://itol.embl.de/). Accessions used are MdHSFA8a (NP_001281294.1), GmHSFB2b (XP_014619265.1), AtHSFA2 (AT2G26150), AtHSF1 (AT4G17750), AtHSF3 (AT5G16820), AtHSFA8 (AT1G67970), AtHSFC1 (AT3G24520), AtHSFA3 (AT5G03720), AtHSF4 (AT4G36990), AtHSFB2A (AT5G62020), AtHSFB2B (AT4G11660), and AtHSFA4A (AT4G18880). Md, *Malus domestica*; Gm, *Glycine max*; At, *Arabidopsis thaliana.* (b) Subcellular localization of MaHSF11 in *N. benthamiana* leaves. The YFP-MaHSF11 fusion protein localizes to both the nucleus and the cytoplasm and coincides with the signal from the nuclear localization signal-red fluorescent protein (NLS-RFP) fusion in *N. benthamiana* leaves as observed in confocal microscopy. Empty vector pSITE-3CA as used as negative control to visualize free YFP. Scale bar, 20Lμm. (c) Relative expression of *MaHSF11* genes in different tissues of banana as determined by RT-qPCR. Values shown are means ± SD of three biological replicates. (d) GUS staining of banana fruit discs indicating successful transformation with the empty vector pANIC6b (EV) and the *proZmUBI:MaHSF11* construct. (e) Relative expression of *MaHSF11*, *MaFLS1*, and *MaFLS2* in transiently transformed banana fruit discs overexpressing (OE) *proZmUBI:MaHSF11* as determined by RT-qPCR. The empty vector was transformed as a control (expression level set to 1). Values shown are means ± SD of three technical replicates of pool of 4-5 independent biological samples. (f) Quantification of the flavonols kaempferol, quercetin, and myricetin in transformed banana fruit discs. (g) GUS staining of banana fruit discs indicating successful transformation with the empty vector EV and the *proZmUBI:MaHSF11*-RNAi construct. (h) Relative expression of *MaHSF11, MaFLS1*, and *MaFLS2* in *MaHSF11*-silenced banana fruit discs transiently expressing *proZmUBI:MaHSF11*-RNAi as determined by RT-qPCR. The empty vector pANIC8b was transformed as a control (expression level set to 1). Values shown are means ± SD of three technical replicates of pool of 4-5 independent biological samples. (i) Quantification of the flavonols kaempferol, quercetin, and myricetin in EV and *proZmUBI:MaHSF11*-RNAi banana fruit discs. Values are represented as means ± SD of three replicates. Unpaired student’s t-tests were used for statistical analyses. *PL≤L0.05; **PL≤L0.01; ***PL≤L0.001.

To further test the regulatory role of MaHSF11, we transiently overexpressed *proZmUBI:MaHSF11* in banana fruit discs (Fig. 3d). RT-qPCR analysis confirmed that the expression of *MaHSF11* was significantly higher (8-fold, *P* ≤ 0.001) in *proZmUBI:MaHSF11* OE banana fruit disc tissues than in the EV controls (Fig. 3e). Furthermore, the expression of *MaFLS1* was significantly upregulated (3-fold, *P* ≤ 0.01), and the expression of *MaFLS2* was downregulated about 2-fold (Fig. 3e). We also observed differential expression of other structural flavonol biosynthesis genes in *MaHSF11*-OE banana fruit tissues (Supplemental Fig. S4a-d) as well as ∼3-fold upregulation in MaMYBFA3 expression, but no significant upregulation of the other two MYB genes (Supplemental Fig. S4e). We quantified the levels of several different flavonol compounds in *MaHSF11*-OE fruit tissues. LC-MS-based quantification showed increased levels of kaempferol, quercetin, and myricetin in transgenic fruit discs (Fig. 3f). Taken together, these results suggest that MaHSF11 positively regulates flavonol synthesis via regulation of several key biosynthesis genes, including direct regulation of *MaFLS1*.

To further verify the regulatory role of MaHSF11 in banana, we transient expressed pANIC8b-EV and *proZmUBI:MaHSF11*-RNAi constructs in pANIC8b in immature banana fruit discs. GUS staining of transformed fruit discs revealed successful integration of the expression cassette in banana (Fig. 3g). Transcript analysis revealed decreased expression of *MaHSF11* in *MaHSF11*-silenced banana fruit tissues suggesting successful gene silencing (Fig. 3h). As expected, the expression level of *MaFLS1* decreased significantly, with the significant increase in the expression of *MaFLS2* (Fig. 3h). Further, transcript analysis of the other biosynthesis genes revealed insignificant changes in the expression level of many biosynthesis genes except a few (Supplemental Fig. S5a-d). Interestingly, we found significant increase in the transcript level of *MaMYBFA1*, but not in *MaMYBFA2* and *MaMYBFA3* genes, in *MaHSF11*-silenced tissues suggesting either direct or indirect negative regulatory role of MaHSF11 over *MaMYBFA1* (Supplemental Fig. S5e). Promoter analysis revealed the absence of any HSE element in the promoter of *MaMYBFA1* indicating no probable direct interaction between the two (Supplemental File S1). Finally, metabolite analysis showed decreased accumulation of kaempferol, quercetin and myricetin derivatives in *MaHSF11* silenced tissues (Fig. 3i). These results further reiterate the regulatory role of MaHSF11 in flavonol biosynthesis in banana.

### MaHSF11 directly regulates *MaF3’5’H1* to promote flavonoid B-ring hydroxylation and enhances myricetin biosynthesis

MaHSF11 overexpression led to significantly higher up-regulation of all the *MaF3*′*5*′*H* gene family members except *MaF3’5’H7* (Fig. 4a). *MaF3*′*5*′*H1* was the most upregulated gene (a 7-fold increase), followed by *MaF3*′*5*′*H*2 and other homologs; *MaF3*′*H* expression decreased negligibly (Fig. 4a, Supplemental Fig. S4d). In *MaHSF11*-silenced tissues, we observed significantly decreased expression in the transcript of *MaF3’H*, *MaF3’5’H4* and *MaF3’5’H5*, while insignificant alteration was observed in the transcript level of other paralogs (Fig. 4b, Supplemental Fig. S5d).

**Figure 4.**
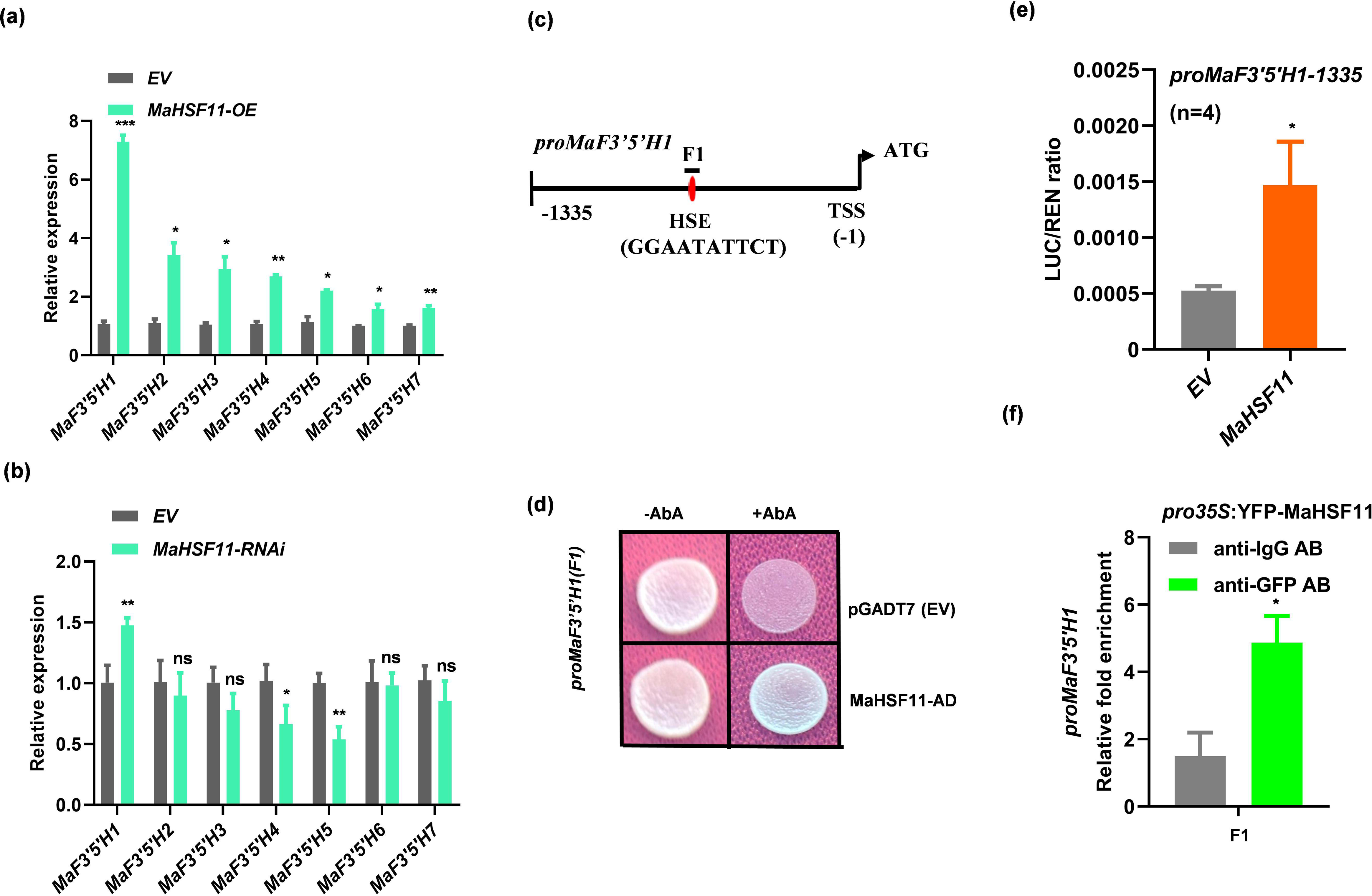
MaHSF11 promotes flavonoid B-ring hydroxylation by modulating the expression of *MaF3*′*5*′*H1*. (a) Relative expression of *MaF3*′*5*′*H1* genes in transformed banana fruit discs transiently overexpressing *proZmUBI:MaHSF11*, as determined by RT-qPCR (EV, empty vector control). Values shown are means ± SD of three technical replicates of pool of 4-5 independent biological samples. (b) Relative expression of *MaF3*′*5*′*H1* genes in *MaHSF11*-silenced banana fruit discs, as determined by RT-qPCR (EV, empty vector control). Values shown are means ± SD of three technical replicates of pool of 4-5 independent biological samples. (c) Illustration of the *cis*-motifs in the region located within 1335 bp upstream of the TSS in the *MaF3’5’H1* promoter. F1 fragment contain one HSE element having GGAATATTCT motifs. (d) Y1H assay showing the interaction of MaHSF11 with *proMaF3*′*5*′*H1–100* fragment F1 carrying HSE (GGAATATTCT). pGADT7 was used as a negative control. Growth was checked on synthetic dropout medium supplemented with AbA. (e) Dual-luciferase assay showing transactivation of *proMaF3*′*5*′*H1–1335* by MaHSF11. The promoter sequence driving the expression of *luciferase* (LUC) was co-infiltrated with the EV or *MaHSF11* constructs in *N. benthamiana* leaves. Values are shown as means ± SD of the LUC/REN ratio of four independent biological replicates (*n* = 4). (f) ChIP-qPCR assay showing binding of MaHSF11 to fragment F1 of *proMaF3*′*5*′*H1* carrying an HSE (nGAAnnTTCn). The promoter fragment carrying HSE was co-infiltrated into *N. benthamiana* leaves with the *pro35S:YFP-MaHSF11* construct. Fold enrichment values are represented as means ± SD of two biological replicates. An anti-IgG antibody was used as a negative control, and an anti-GFP antibody was used to precipitate MaHSF11-enriched chromatin. Unpaired Student’s *t*-tests were used for statistical analyses. *\*P*□≤□0.05.

Thorough analysis of the promoter sequences of these genes revealed the presence of one or more HSE elements in the promoters of all these genes except *MaF3’5’H7*, suggesting direct regulation by MaHSF11 (Supplemental File S1). The more than 7-fold up-regulation of *MaF3’5’H1*, (the highest among all the paralogs) prompted us to check whether it can be directly regulated by MaHSF11 at the transcriptional level. Promoter of *MaF3*′*5*′*H1* contains a single HSE (nGAAnnTTCn) at the –720-bp position relative to the TSS (Fig. 4c, Supplemental File S1). We performed Y1H assays to test the interaction of MaHSF11 with the HSE of *proMaF3*′*5*′*H1–1335* using a 100-bp, HSE-containing promoter fragment (F1), which showed a strong interaction with MaHSF11, as indicated by yeast growth (Fig. 4c,d). The ability of MaHSF11 to transactivate *proMaF3*′*5*′*H1–1335* was tested in a dual-luciferase assay in *N. benthamiana* leaves. The increased LUC activity with MaHSF11, compared to the EV control, indicated that MaHSF11 activated the *MaF3*′*5*′*H1* promoter (Fig. 4e). To further test the direct binding of MaHSF11 via the HSE, we co-infiltrated a *pro35S:YFP-MaHSF11* construct and the *proMaF3*′*5*′*H1–1335* fragment into *N. benthamiana* leaves and performed a ChIP assay using an anti-GFP antibody. This ChIP-qPCR assay revealed enrichment of the HSE-containing 100-bp promoter fragment F1 (Fig. 4f). Overall, these results show that MaHSF11 directly regulates the expression of *MaF3*′*5*′*H1*.

### MaHSF11 regulates *MaMYBFA3* through MaDREB1

Overexpression and gene silencing of *MaHSF11* in banana fruit discs resulted in significant alteration in the expression level of several other structural genes involved in flavonol biosynthesis (Supplemental Fig. S4a-d, Supplemental Fig. S5a-d). Therefore, we investigated the expression of regulatory *MaMYBFA1–3* genes. As stated earlier, we found a significant upregulation of only *MaMYBFA3* in banana fruit discs overexpressing *MaHSF11* (Supplemental Fig. S4e), while no significant change was observed in *MaHSF11* silenced tissues (Supplemental Fig. S5e). A promoter analysis of the ∼2-kb region upstream of the *MaMYBFA3* TSS revealed the absence of any HSEs (Supplemental File S1). Therefore, we hypothesized that MaHSF11 might indirectly regulate the expression of *MaMYBFA3* via an unknown factor. Furthermore, promoter analysis of *MaMYBFA3* revealed the presence of several DRE/CRT (CCGAC) elements in the ∼2.5-kb region upstream of the TSS (Supplemental File S1). Because HSFs bind HSEs in the promoter of *DREB* genes (Yoshida *et al*., 2011; Huang *et al*., 2016), we hypothesized that DREB TFs may act as mediators between *MaHSF11* and *MaMYBFA3*. To investigate the likelihood that MaHSF11– MaDREB–MaMYBFA3 is a regulatory module in banana, we measured the expression levels of *DREB* genes using the RNA-Seq data from drought-treated bananas of the Grand Naine cultivar (Muthusamy *et al*., 2016). We found the expression of a single *DREB* gene (*Ma04_t2630*), which was also upregulated after drought stress (Fig. 5a). This gene was previously characterized as MaCBF1 (C-repeat binding factor1) and described as a critical regulator of cold stress tolerance in banana (Zhao *et al*., 2013; Shan *et al*., 2014). We isolated the 777-bp full-length CDS of this gene and named it *MaDREB1* due to nucleotide sequence dissimilarity with MaCBF1 (Shan *et al*., 2014). Phylogenetic analysis revealed that MaDREB1 formed a clade with rice OsDREB3 and maize ZmDREB (Supplemental Fig. S6a). Further, subcellular localization assay in *N. benthamiana* leaves using a pro35S:YFP-MaDREB1 construct confirmed nuclear localization of YFP-MaDREB1, which co-localized with a nuclear NLS-RFP marker (Supplemental Fig. S6b). In the tissue specific transcript accumulation profiling of *MaDREB1*, we found that it followed the same distribution pattern as *MaHSF11*, with higher expression level in root, leaf, and least expression in ripe peel and pulp tissue (Supplemental Fig. S6c). To examine whether the promoter of *MaDREB1* was regulated by MaHSF11, we analyzed the region ∼2 kb upstream of the TSS and identified two HSEs at positions –223 and –576 bp upstream of the TSS (Supplemental File S1). We measured the expression of *MaDREB1* in *MaHSF11* OE fruit discs and determined that it was significantly upregulated (>6-fold) (Fig. 5b). Analysis of *MaDREB1* expression in *MaHSF11* silenced fruit tissues showed insignificant alteration in the expression level of *MaDREB1* (Fig. 5c)

**Figure 5.**
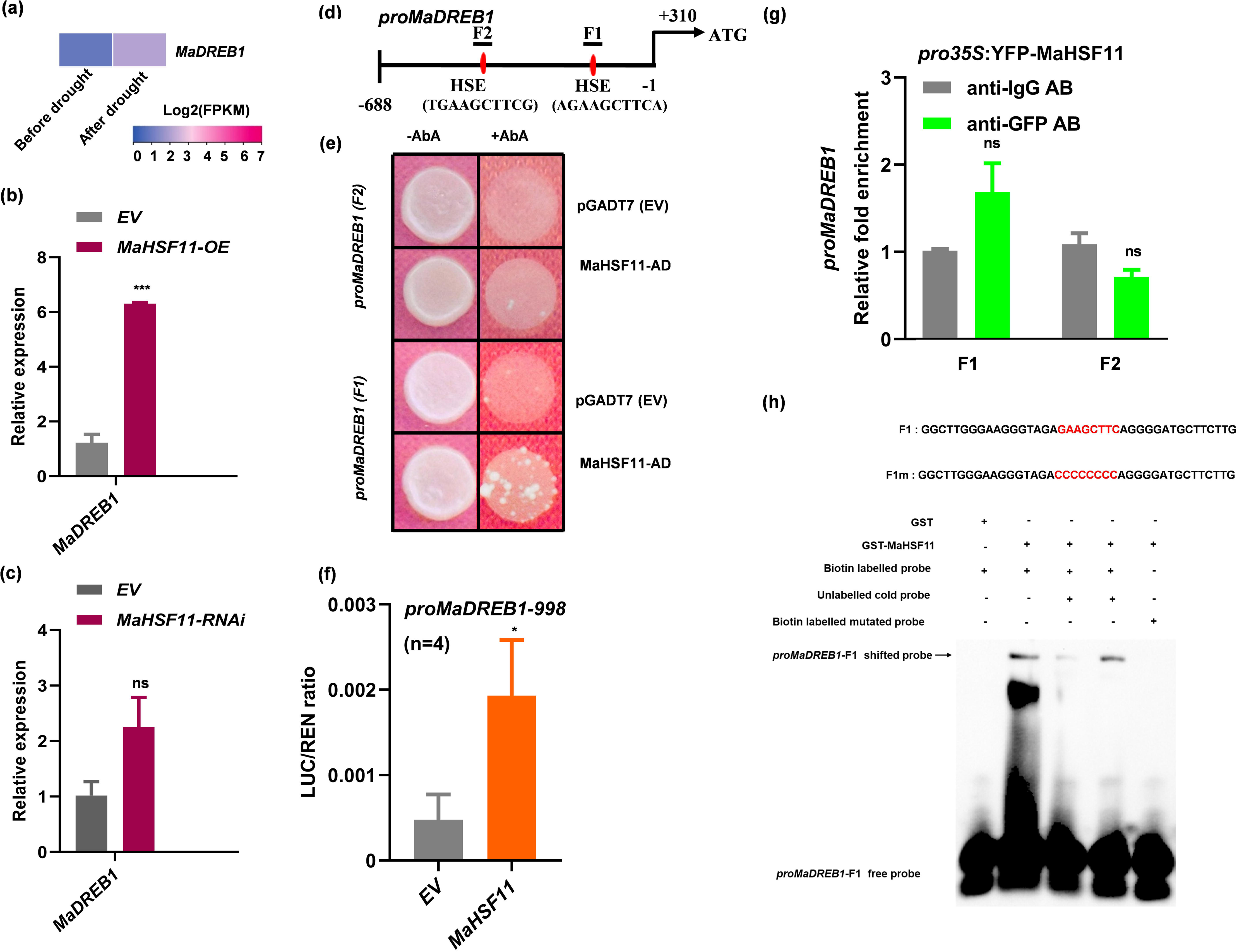
MaHSF11 acts upstream of *MaDREB1* to positively regulate its expression in banana. (a) Heatmap showing expression of candidate *MaDREB* genes before and after drought treatment in banana (Grand Naine cultivar, AAA genome). FPKM values for the indicated genes were extracted from publicly available RNA-Seq datasets and plotted according to the 0–7 color scale shown. (b) Relative expression of *MaDREB1* in banana fruit discs transiently transformed with EV or *proZmUBI:MaHSF11* as determined by RT-qPCR. (c) Relative expression of *MaDREB1* in *MaHSF11*-silenced banana fruit discs and EV control as determined by RT-qPCR. (d) Illustration of the *cis*-motifs in the region located within 688 bp upstream of the TSS in the *MaDREB1* promoter. F1 and F2 fragments contain one HSE element each having AGAAGCTTCA and TGAAGCTTCG motifs. +310 shows the bp distance between TSS and start codon. (e) Y1H assay showing interaction of MaHSF11 with the F1 and F2 fragments of *proMaDREB1–100* carrying an HSE (nGAAnnTTCn). pGADT7 was used as a negative control. Growth was checked on synthetic dropout medium supplemented with AbA. (f) Dual-luciferase assay showing transactivation of *proMaDREB1– 998* by MaHSF11. Promoter sequence driving the expression of *LUC* was co-infiltrated with empty vector and *MaHSF11* construct in *N. benthamiana* leaves. Values shown are means ± SD of LUC/REN ratio (*n* = 4 independent biological replicates). (g) ChIP-qPCR assay showing binding of MaHSF11 to *proMaDREB1*. A promoter fragment carrying HSE was co-infiltrated into *N. benthamiana* leaf cells with the *pro35S:MaHSF11-YFP* construct. Fold enrichment values are represented as means ± SD of two biological replicates. An anti-IgG antibody was used as a negative control, and an anti-GFP antibody was used to precipitate MaHSF11 enriched chromatin. Unpaired Student’s t-tests were used for statistical analyses. *\*P*□≤□0.05; *\*\*\*P*□≤□0.001; ns, not significant. (h) EMSA showing the binding of MaHSF11-GST protein with the F1 fragment of *MaDREB1* promoter (*MaDREB1*-F1) *in vitro*. Binding cis-motif and the mutated motif in the probes are highlighted in red in F1 and F1m, respectively. MaHSF11-GST protein, labelled and unlabeled probes were separated in a polyacrylamide gel, blotted on nylon membrane and detected by chemiluminescence. + indicates presence and – indicates absence of probe/protein.

To verify the direct regulation of MaDREB1, we conducted a Y1H assay. We cloned two 100-bp fragments (F1 and F2) harboring HSEs (HSE1 and HSE2, respectively) of the *proMaDREB1* and conducted a Y1H assay with MaHSF11 as the effector (Fig. 5d-e). Promoter fragments carrying the HSEs were cloned into the pAbAi vector and transformed into yeast (*Saccharomyces cerevisiae*) Y1H Gold strain together with pGADT7 as a negative control. MaHSF11 in pGADT7 was transformed with both fragments, and the growth assay was performed on media supplemented with aureobasidin A (AbA). MaHSF11 could bind the HSE1-containing fragment F1 of *proMaDREB1 in vitro* (Fig. 5e). To test the transactivation activity of MaHSF11 on *proMaDREB1*, we cloned a 998-bp promoter fragment of *MaDREB1* into the p635nRRF vector and performed dual-luciferase assays. The EV as a negative control and *MaHSF11* as the effector were individually infiltrated together with the reporter construct carrying a particular promoter fragment into *N. benthamiana* leaves, and the assay was performed 3 days post infiltration. LUC activity increased significantly when MaHSF11 was co-transfected compared to the EV control (Fig. 5f). Furthermore, *pro35S:YFP-MaHSF11* and the promoter fragment were also transformed into *N. benthamiana* leaves, and a ChIP assay was conducted. An anti-GFP antibody was used to precipitate MaHSF11-bound chromatin, and anti-IgG was used as a negative control. ChIP-qPCR showed enrichment of the F1 fragment in the immunoprecipitated chromatin, demonstrating that MaHSF11 directly bound the F1 fragment and regulated *MaDREB1* (Fig. 5g). In addition, EMSA showed direct binding of MaHSF11-GST with the GAAGCTTC motif containing promoter fragment F1 of *MaDREB1* promoter *in vitro* (Fig. 5h). Taken together, these findings strongly suggests that MaHSF11 directly binds to the promoter of *MaDREB1* to enhance its expression.

### MaDREB1 directly regulates the expression of *MaMYBFA3* in banana

To test whether MaDREB1 acted as a direct transcriptional activator of *MaMYBFA3*, we isolated *proMaMYBFA3–1892*, which contains all CRT/DREs (CCGAC/GTCGG), and cloned it into pAbAi. We performed a Y1H assay with MaDREB1 as the effector in pGADT7 and EV as a control. Promoter fragments cloned into the pAbAi vector were transformed into the Y1H strain, and a growth assay was performed on AbA-supplemented media. Growth was observed in the MaDREB1 transformed strain, suggesting that it bound to the promoter fragment (Fig. 6a). To study the transactivation activity of MaDREB1, we cloned a 2508-bp fragment of *proMaMYBFA3* into p635nRRF to perform a dual-luciferase assay with MaDREB1 in pBTDest as the effector and the EV as a negative control. LUC activity increased when MaDREB1 was used as the effector compared to the control, indicating its ability to transactivate *proMaMYBFA3* (Fig. 6b). Overall, these findings show direct regulation of *MaMYBFA3* by MaDREB1. To validate the direct binding of the MaDREB1 on to the promoter of MaMYBFA3 *in vitro*, we conducted EMSA assay with four different probes (F1-F4) having different DRE/CRT motifs designed from the promoter of MaMYBFA3 (Fig. 6c). The EMSA result demonstrated the binding of MaDREB1-GST with the promoter fragment F3 having GTCGG motif, but not with promoter fragment F1, F2, and F4 of the MaMYBFA3 promoter despite having CCGAC/GTCGG motif (Fig. 6d). These results suggests that MaDREB1 specifically binds to F3 region of the MaMYBFA3 promoter to enhance its expression.

**Figure 6.**
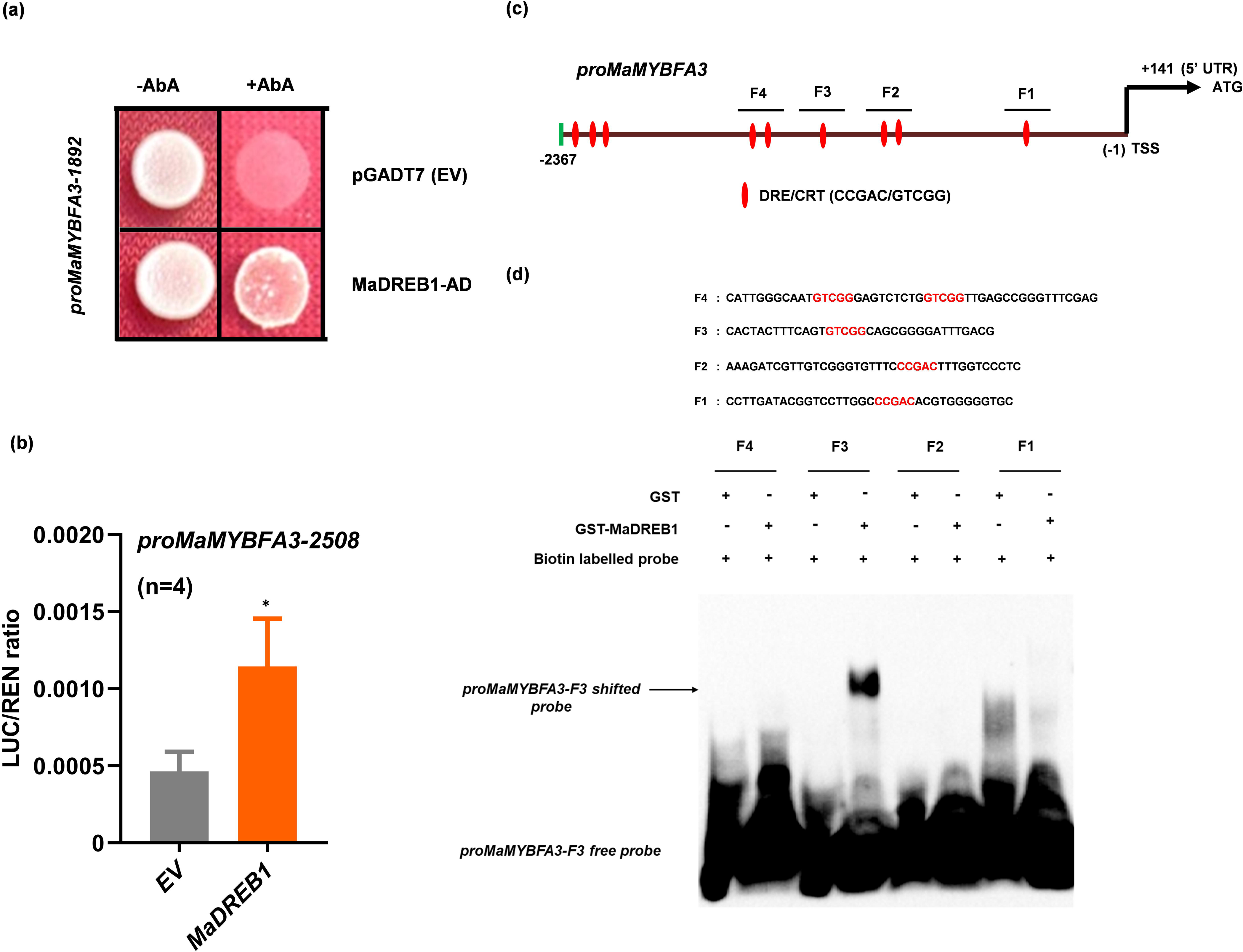
MaDREB1 directly binds to the promoter of *MaMYBFA3* to regulate flavonol biosynthesis. (a) Y1H assay showing interaction of MaDREB1 with the *proMaMYBFA3– 1892* fragment carrying CRT/DREs (CCGAC/GTCGG). pGADT7 was used as a negative control. Growth was checked on synthetic dropout medium supplemented with AbA. (b) Dual-luciferase assay showing transactivation of *proMaMYBFA3–2508* by MaDREB1. The *proMaMYBFA3–2508* sequence driving the expression of *LUC* was co-infiltrated with the empty vector or *MaDREB1* construct into *N. benthamiana* leaves. Values are shown as means ± SD of the LUC/REN ratio (*n* = 4 independent biological replicates). (c) Illustration of the cis-motifs in the region located within 2367 bp upstream of the TSS in the *MaMYBFA3* promoter. F1, F2, F3 and F4 fragments contain one to two DRE/CRT elements having either CCGAC or GTCGG motifs. +141 shows the bp distance between TSS and start codon. (d) EMSA showing the binding of MaDREB1-GST protein with the F3 fragment of *MaDREB1* promoter (*proMaDREB1-F3*) *in vitro*. Different cis-motifs (CCGAC/GTCGG) present in the F1, F2, F3 and F4 probes are highlighted in red. MaDREB1-GST protein, labelled and unlabeled probes were separated in a polyacrylamide gel, blotted on nylon membrane and detected by chemiluminescence. + indicates presence and – indicates absence of probe/protein.

To verify the function of *MaDREB1 in planta*, we transiently overexpressed it in banana fruit discs. GUS staining indicated successful transformation of the GUS expression cassette (Fig. 7a). RT-qPCR analysis of the transformed fruit disc tissue showed high expression (∼375-fold, *P* < 0.0001) of *MaDREB1* relative to the EV control, demonstrating successful overexpression (Fig. 7b). Expression analysis of *MaMYBFA3* in the *MaDREB1* OE tissue showed an ∼10-fold increase in the expression level (*P* < 0.05). We also found higher expression (∼21-fold, *P* < 0.001) of *MaMYBFA2*, while *MaMYBFA1* expression was unaltered (Fig. 7c). *MaFLS1* and *MaFLS2* also showed opposite trends in expression in the *proZmUBI:MaDREB1* OE fruit tissues: *MaFLS1* was downregulated, and *MaFLS2* was upregulated (Fig. 7c). Differential expression of other structural flavonol biosynthesis–related genes was also observed in *proZmUBI:MaDREB1* OE banana fruit tissues (Supplemental Fig. S7a-e). To study the modulation of flavonol content resulting from *MaDREB1* overexpression, we used LC-MS to quantify the levels of different metabolites and found significantly higher accumulation of kaempferol, quercetin, and myricetin (Fig. 7d). These results indicate that MaDREB1 positively regulates flavonol biosynthesis in banana.

**Figure 7.**
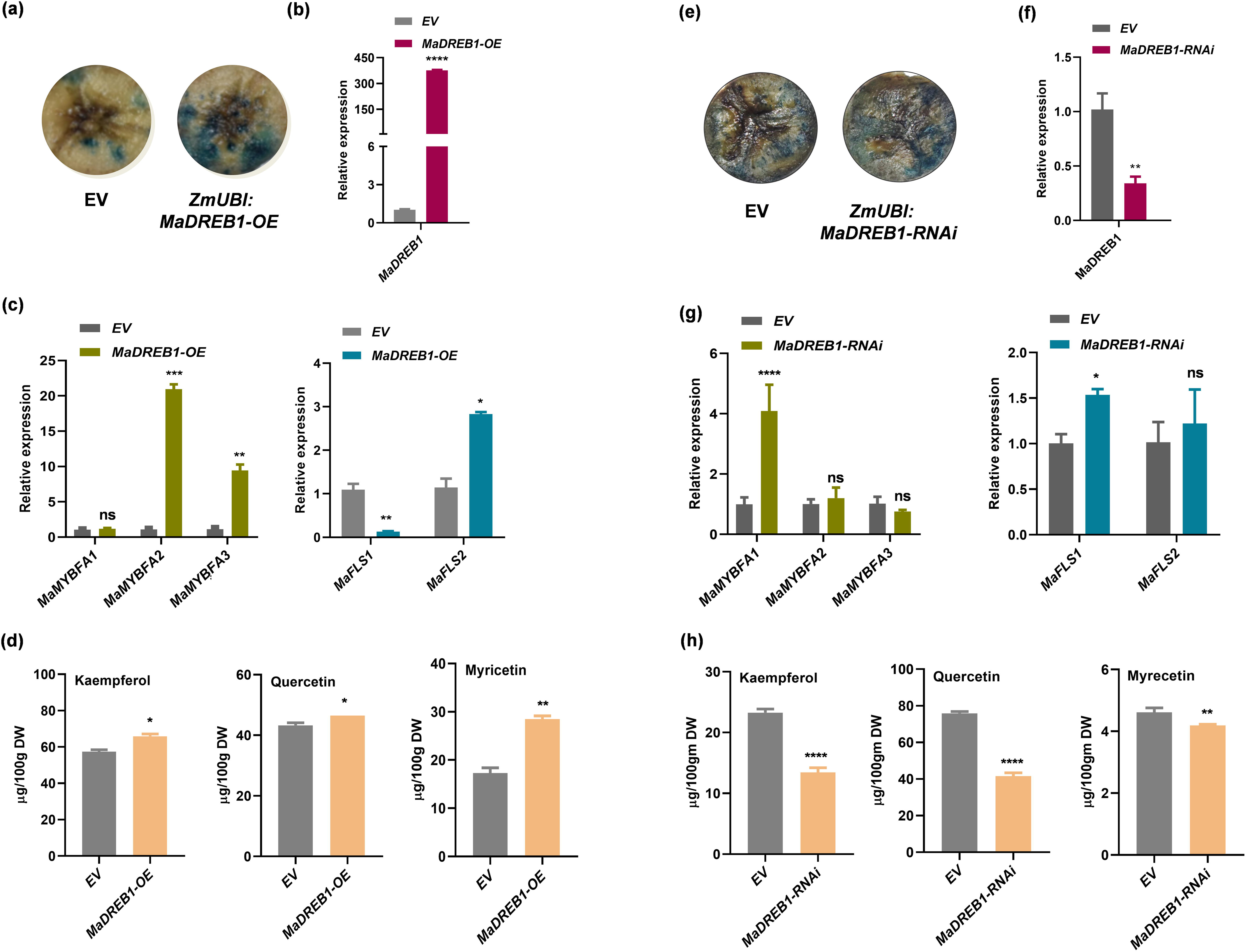
MaDREB1 regulates the expression of *MaMYBFAs*, *MaFLS1*, and *MaFLS2* to modulate flavonol biosynthesis in banana. (a) GUS staining of EV and *proZmUBI:MaDREB1* transformed fruit discs. (b) RT-qPCR-based relative expression of *MaDREB1*, and (c) relative expression of *MaMYBFAs*, *MaFLS1*, and *MaFLS2* in transformed banana fruit discs transiently expressing EV and *proZmUBI:MaDREB1*. EV was transformed as a control. Values shown are means ± SD of three technical replicates of a pool of 4-5 biological replicates. (d) Quantification of the flavonols kaempferol, quercetin, and myricetin in transformed fruit discs. (e) GUS staining of EV and *proZmUBI:MaDREB1-RNAi* transformed fruit discs. (f) RT-qPCR-based relative expression of *MaDREB1, and (g)* relative expression of *MaMYBFAs, MaFLS1*, and *MaFLS2* in *MaDREB1*-silenced banana fruit discs transiently expressing *proZmUBI:MaDREB1-RNAi* and EV. EV pANIC8b was transformed as a control. Values shown are means ± SD of three technical replicates of pool of 4-5 independent biological pools. (h) Quantification of the flavonols kaempferol, quercetin, and myricetin in transiently expressing *proZmUBI:MaDREB1*-RNAi banana fruit discs. Values are represented as means ± SD of three replicates. Unpaired Student’s t-tests were used for statistical analyses. *PL≤L0.05; **PL≤L0.01; ***PL≤L0.001; ****P ≤ 0.0001.

To further verify the regulatory role of MaDREB1 in banana, we did gene silencing of MaDREB1 in immature banana fruit. GUS staining in pANIC8b and *proZmUBI:MaDREB1*-RNAi expressing banana fruits revealed successful integration of the expression cassette in banana (Fig. 7e). Expression analysis showed decreased transcript of *MaDREB1* in *MaDREB1*-silenced banana disc fruit tissues suggesting successful gene silencing (Fig. 7f). We did not see any significant changes in the expression level of *MaMYBFA2* and *FA3* (Fig. 7g). Surprisingly, the transcript level of *MaMYBFA1* increased by 4-fold suggesting that the later might be directly or indirectly regulated by MaDREB1 (Fig. 7g). As expected, the expression level of *MaFLS1* opposite trends as compared to overexpression, it increased significantly. We did not observe any significant change in the transcript of *MaFLS2* (Fig. 7g). Transcript analysis of the other biosynthesis genes revealed significant increase in the expression level of many biosynthesis genes, except few genes where no change was observed (Supplemental Fig. S8a-e). Finally, metabolite analysis showed decreased accumulation of kaempferol, quercetin and myricetin derivatives in *MaDREB1* silenced tissues (Fig. 7h). Spatio-temporal metabolite accumulation is also controlled at post-translational level apart from transcriptional regulation. These results reveled that MaDREB1 has an overall regulatory role to modulate flavonol biosynthesis in banana.

### Heat stress positively modulates flavonol biosynthesis in banana

HSFs, as the terminal component of the heat stress response pathway, also regulate the expression of many thermo-responsive genes (von Koskull-Döring *et al*., 2007; Liu *et al*., 2013). Flavonol synthesis is modulated by heat stress conditions, wherein flavonols are known to scavenge excess ROS (Muhlemann *et al*., 2018). Since MaHSF11 was induced under drought stress (Fig. 2a), we wanted to check whether the expression level of *MaHSF11* and the flavonol biosynthesis pathway are modulated upon exposure to heat stress. To do this, we exposed three-month-old banana plantlets to 42°C under optimal photoperiod and light conditions, while control plants were kept at 26°C (Fig. 8a). Expression analysis revealed increased transcript levels of *MaHSF11* at high temperature as compared to the control plants (Fig. 8b). Similarly, we found increased expression levels of *MaDREB1*, *MaMYBFA1*-3, *MaFLS1*, and *MaFLS2* under heat stress conditions compared to control, each at different time points (Fig. 8b). We further checked the expression of other biosynthesis genes and found that transcript levels of a few genes up-regulated while a few genes showed down-regulation upon heat stress (Supplemental Fig. S9). Interestingly, different paralogs of *MaF3’5’H* genes showed increased transcript levels under heat stress at different time points (Supplemental Fig. S9). Finally, metabolite analysis showed modulation in the accumulation of kaempferol, quercetin, and myricetin derivatives under heat stress (Fig. 8c). Quercetin content decreased at early time points, while kaempferol content fluctuated at different time points (Fig. 8c). However, we observed a sharp increase in the accumulation of myricetin right after the heat treatments, gradually up to 24 hours (Fig. 8c), which may be attributed to the up-regulation of different *MaF3’5’H* paralogs at different time points of heat stress treatment. Taken together, our results show that flavonol biosynthesis is responsive to heat stress conditions in banana, and that *MaHSF11* might also contribute to enhanced flavonol accumulation under heat stress in banana.

**Figure 8.**
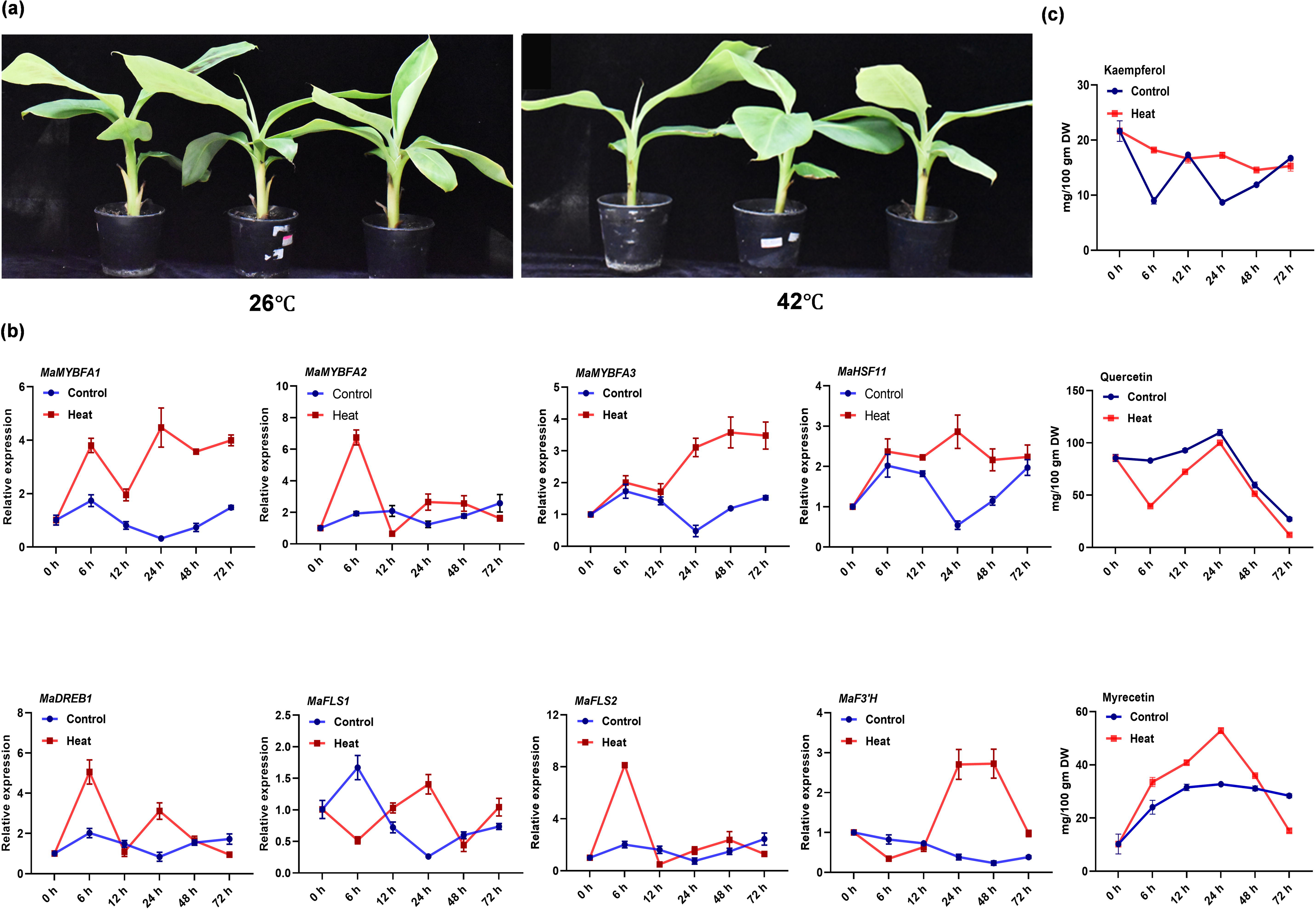
Heat stress positively modulates flavonol biosynthesis in banana. (a) Representative images of three months old banana seedlings exposed heat stress at 42L. Control seedlings were kept at 26L. (b) RT-qPCR-based relative expression of *MaHSF11, MaDREB1, MaMYBFA1-3, MaFLS1* and *MaFLS2* at different time points (0h, 6h, 12h, 24h, 48h, 72h) of heat stress (42L) treatment in banana seedlings, and control (26L) seedlings. (c) Quantification of the flavonols aglycones i.e., kaempferol, quercetin, and myricetin in the heat stress treated and control plants at different time intervals. Values are represented as mean ±SD of three biological replicates.

## Discussion

Flavonoid biosynthesis pathways developed during the evolution of land plants. As they became terrestrial, plants developed response mechanisms to adapt to and cope with several new environmental conditions. Flavonoid biosynthesis is highly responsive to drought stress along with other biotic and abiotic stress conditions (Wang *et al*., 2020; Wang *et al*., 2021b; Naik *et al*., 2022; Song *et al*., 2022; Gautam *et al*., 2023). The role of stress signalling in the context of flavonoid biosynthesis has been studied recently, strengthening our knowledge regarding transcriptional regulation of flavonoid biosynthesis. Although flavonol biosynthesis, in particular, is positively modulated by abiotic and biotic stress, a conserved response seen throughout the plant kingdom (Albert *et al*., 2018; Naik *et al*., 2022; Davies *et al*., 2023; Lu *et al*., 2023), the molecular mechanism controlling this response has remained obscure. In this study, we identified a novel transcriptional regulatory network involving different classes of TFs and showed their interaction in regulating SG7 *MaMYBFA* and flavonol biosynthesis genes in the herbaceous fruit crop banana. We deciphered important details of the molecular mechanisms involving the transcriptional regulatory module of how HSFs, DREB, and MYB proteins orchestrate flavonol accumulation in abiotic stress dependent manner.

Initially, three candidate SG7 R2R3-MYB TFs, *MaMYBFA1–3*, were identified as having high sequence conservation in their R2R3 domains and partial conservation in the SG7-1 (GRTxRSxMK, Stracke *et al*., 2001) and SG7-2 motifs ([W/x] [L/x]LS, Czemmel *et al*., 2009). Furthermore, the absence of the bHLH-interacting motif ([DE]Lx2[RK]x3Lx6Lx3R, Zimmermann *et al*., 2004) in the R3 domain suggested that these three candidate genes were independent regulators of flavonol synthesis in banana. Expression analysis of the three *MaMYBFA* genes revealed the highest level of transcript accumulation in leaf and bract tissue. *MaFLS1* transcript level was highest in bract and leaf tissues as compared to other tissues, whereas *MaFLS2* showed highest expression in bract tissue only. Different flavonol derivatives, on the contrary, are accumulated varyingly in different tissues, suggesting multi-layered regulation of the metabolic pathway. This finding indicates the presence of a regulator other than SG7 MYB in banana, acting upstream of *MaFLS1*. All these results prompted us to functionally characterize the candidate *MaMYBFA* genes in banana to determine their possible roles as flavonol-specific TFs. To do this, we transiently overexpressed the *MaMYBFA* genes in banana fruit. As raising stable transgenics in banana is time-consuming process which takes years of laborious efforts, we preferred transgene expression system in immature banana fruit as it is well established and has been used in many studies (Matsumoto *et al*., 2009; Xiao *et al*., 2018: Wu *et al*., 2019; Wei *et al*., 2023a). Although *MaFLS2* expression was activated by MaMYBFA, the expression of *MaFLS1* was downregulated. SG7 R2R3 MYB TFs directly regulates the expression of *FLS* genes in other species. Thus, *MaFLS1* also seems a direct target of MaMYBFAs in banana. If it is true, then later might depend on some associative or interacting partner for strong repression. Few TFs have been reported to directly repress the expression of *FLS* genes in other species (Zhang *et al*., 2018; Bian *et al*., 2020; Wang *et al*., 2021d). It is also apparent that binding is mediated by MaMYBFAs, as we did not find any known repression motifs in the protein sequences of the concerned. The repression may be affected by the interacting partner, which might be spatio-temporally regulated in banana. Few instances have also been documented in previous studies, where *FLS2* genes are strongly repressed by transcription regulators. In *Fagopyrum tataricum*, the expression level of *FtFLS* and rutin synthesis is severely affected in *FtMYB13*, *FtMYB14*, *FtMYB15*, *FtMYB16* and *FtJAZ1* overexpressing hairy roots (Zhang *et al*., 2018). FtMYB13-16, the four SG4 R2R3 MYBs, directly interacts with FtJAZ1, suggesting that *FtFLS1* expression is repressed by cooperative action of both the factors. *MdBEH2*.2 directly binds to the E-box region in the promoter of *MdFLS* to repress the expression (Wang *et al*., 2021d). It also physically interacts with MdMYB60 and overexpression of *MdMYB60* in apple calli leads to down-regulation of *MdFLS*. GmNAC2 directly binds to and represses the expression of *GmFLS1* and *GmFLS2* in soybean (Bian *et al*., 2020). These reports hints at the role of probable interacting partners of MaMYBFA, which may influence the transcriptional activity of MaMYBFA strongly. On the other hand, DELLAs (RGA and GAI) directly interacts with MYB12 and MYB111 to enhance the transcriptional activation activity of the later over the promoter of *FLS* in Arabidopsis (Tan *et al*., 2019). In *Camellia sinensis*, CsMYB12 interacts physically with CsbZIP1, and both directly binds to promoter of *CsFLS* to promote flavonol synthesis (Zhao *et al*., 2021). In an antagonistic way, CsMYB4 and CsMYB7 directly repressed the expression of *CsFLS*. They also interacted with CsMYB12 to attenuate the transcriptional activation of the later (Zhao *et al*., 2021). All these reports suggest that some strong repressors might be operating in banana in the same way as CsMYB4 and CsMYB7. This finding could be an exception, because none of the SG7 MYB activators downregulate flavonol synthase genes, the key genes in the flavonol biosynthesis pathway. Flavonol content mainly remained unaltered with MaMYBFA overexpression, a finding that can be attributed to the downregulation of *MaFLS1* and *MaF3*′*H* and increased expression of *MaFLS2*, which could compensate for it. Thus, although we concluded that MaMYBFAs are weak regulators of flavonol synthesis in banana, it is difficult to ascertain their positive regulatory role on flavonol synthesis. However, our results show the division of labor among the SG7 MYB transcriptional regulators within the regulatory cascade. Such a mechanism has been observed in the dicot freesia (*Freesia hybrida*), in which four SG7 MYB proteins, FhMYBF1–4, differentially regulate the expression of *FhCHS* and *FhCHI* paralogs, as well as *FhFLS1*, but not *FhFLS2* (Shan *et al*., 2020). SG7 MYB TFs regulate flavonol synthesis by directly binding to MYB binding sites (MBSs, CNGTTR) present in the promoters of flavonol biosynthesis genes, activating their expression. In apple, MdMYB22 binds to the MBS in the promoters of *MdCHS* and *MdFLS* to regulate their expression (Wang *et al*., 2017). In freesia, FhMYB1–4 bind to the MYB core (CNGTTR) and MYBPLANT motifs (MACCWAMC), which are essential for promoter binding and transactivation (Shan *et al*., 2022). Overall, these results suggest that MaMYBFA1–3 are the functional flavonol-specific transcriptional regulators in banana. As many early biosynthesis genes were highly up-regulated upon overexpression of MaMYBFAs, it can be speculated that the MaMYBFAs probably enhances flux towards the overall flavonoid biosynthesis pathway, rather specific flavonol branch. A recent study reported a very interesting rather unusual mechanism in Jojoba, where a flavonol regulator ScMYB12 physically interacts with ScTT8 to regulate the expression of *ScFLS* gene (Zheng *et al*., 2024). SG7 R2R3 MYBs NtMYB12a, apart from their classical role in the regulation of flavonol synthesis, are also reported to regulate fatty acid synthesis in tobacco (Wang *et al*., 2021e). It would be interesting to explore the impact of MaMYBFA on the regulation of other metabolic or developmental pathways in banana.

HSFs are critical components of the signaling cascades involved in responses to salt, cold, drought, and heat stress (Chauhan *et al*., 2011; Bian *et al*., 2020; Wang *et al*., 2020). Several HSFs and their regulatory networks have been discovered in Arabidopsis, rice, tomato, soybean (*Glycine max*), wheat, and grape (*Vitis vinifera*) (Guo *et al*., 2016; Wu *et al*., 2022). In this study, MaHSF11 was identified as a regulator of *MaFLS1*. MdHSFA8a was previously identified as a regulator of *MdFLS* in apple, promoting flavonol biosynthesis and the drought stress response by regulating *MdFLS* expression (Wang *et al*., 2020). GmHSFB2B confers salt stress tolerance and flavonol synthesis in soybean by repressing *GmNAC2*, a negative regulator of *GmFLS1* and *GmFLS2* (Bian *et al*., 2020). Apart from these, other non-SG7 MYB regulators from other species include AtMYB21, AtMYB24, and FhMYB21L2. The presence of three HSEs in the promoter of *MaFLS1* suggested it might be regulated by a HSF. We determined the role of MaHSF11 to be a regulator of CYP450 B-ring hydroxylase in banana, where it directly bound to the promoter of *MaF3*′*5*′*H1* (Fig. 4d). Very few regulators of *F3*′*5*′*H* have been identified in plants. In poplar (*Populus tremula×tremuloides*), MYB117 specifically regulates *F3*′*5*′*H* expression to promote anthocyanin synthesis and flavonoid B-ring hydroxylation (Ma *et al*., 2021). MYBC1 and WRKY44 both independently and cooperatively regulate *F3*′*5’*′*H* expression to regulate the anthocyanin level in kiwifruit (*Actinidia purpurea*) (Peng *et al*., 2020).

In our study, *MaHSF11* overexpression upregulated *MaFLS1* but downregulated *MaFLS2*, indicating the presence of a complex regulatory cascade leading to preference of one paralog over the other. MaHSF11 also promoted the expression of several *MaF3*′*5*′*H* gene homologs, while slightly repressing *MaF3*′*H* (Supplemental Fig. S4d). Since the enzymes compete with each other for the common substrate dihydrokaempferol, MaHSF11 might be specifically promoting the myricetin synthesis branch and tries to suppresses the quercetin synthesis branch catalyzed by MaF3′H, either directly or indirectly. *MaFLS1* was highly expressed in young leaves, and this finding can be correlated with higher *MaHSF11* expression in leaves compared to fruits and roots (Wei *et al*., 2016). Fruit tissue, particularly peel, had higher expression of *MaF3*′*H*, leading to a greater accumulation of rutin and quercetin in peel compared to other tissues (Pandey *et al*., 2016). Of the seven paralogs of *MaF3*′*5*′*H* in banana, five were upregulated 2-fold with *MaHSF11* overexpression, suggesting a role for *MaHSF11* as a strong regulator of this gene family (Fig. 4a). These results further suggest that MaHSF11 promotes flavonol biosynthesis via a distinct regulatory module to regulate stress-responsive genes such as flavonol biosynthesis–related genes involved in drought, cold, and salinity stress responses in banana.

Preference of MaHSF11 for the *MaFLS1* and *MaF3*′*5*′*H* homologs (relative to *MaFLS2* and *MaF3*′*H*) suggested a specificity of MaFLS1 for dihydromyricetin as a substrate. In a recent study, MaFLS1 also showed specificities for dihydrokaempferol and dihydroquercetin. In future studies, it will be important to determine whether MaFLS1 has a substrate specificity for dihydromyricetin as well. CYP450 enzymes are involved in the flavonoid metabolon in both rice and soybean (Shih *et al*., 2008; Waki *et al*., 2016), where they act as ER membrane–bound nucleation sites for metabolon formation. Interaction among enzymes varies greatly in terms of their stability and complexity. Preferential upregulation of MaF3′5′H and MaFLS1 and downregulation of MaF3′H and MaFLS2 suggested probable metabolon formation in the ER, which can be investigated in future. In snapdragon (*Antirrhinum majus*), F3′H associates with CHI to form a metabolon (Fujino *et al*., 2018). In Arabidopsis, FLS1 interacts with CHS via competition with DFR (dihydroflavonol 4-reductase), which can also bind to CHS (Crosby *et al*., 2011). Negative regulation of MaFLS2 by MaHSF11 also strongly suggests functional promiscuity of the MaFLS2 enzyme as it has not been functional characterized. The absence of HSE element in the promoter of MaFLS2 indicates its indirect repression by MaHSF11. In soybean, GmHSFB2b represses the expression of *GmNAC2*, negative regulators of GmFLS1/2. So, two possible mechanisms behind the down-regulation of MaFLS2 can be the up-regulation of any of its repressors or down-regulation of its positive regulators by MaHSF11.

We also observed that MaHSF11 upregulated *MaMYBFA3* indirectly via MaDREB1. Interactions of DREB and HSF TFs have been reported previously (Yoshida *et al*., 2011;Chen *et al*., 2016). In Arabidopsis, DREB2A is transcriptionally regulated by HSFA1a, HSFA1b, and HSFA1d (Yoshida *et al*., 2011), and DREB2A also acts upstream of HSFA3 to regulate its expression (Schramm *et al*., 2008). In our study, we found ∼2-fold increased expression of MaHSF11 when *MaDREB1* was overexpressed in banana, whereas suggesting conservation of an interconnected regulatory cascade (Supplemental Fig. S10). We also determined that MaHSF11 directly bound to the HSE in *proMaDREB1* to regulate its expression. Overexpression of *MaDREB1* significantly down-regulated several biosynthesis genes related to flavonol biosynthesis, while few biosynthesis genes were also up-regulated notably *MaCHS4* and *MaF3’5’H3*. This antagonistic modulation in the expression level of different biosynthesis genes leads to increased flavonol accumulation, although inadequately (Supplemental Fig. S7a–e). While, gene silencing led to significantly up-regulation of few genes (Supplemental Fig. S8a-e). In future work, it will be intriguing to determine whether MaDREB1 directly represses these genes. HSF and DREB are not reported so far to interact physically. In our Y2H analysis, we could not see any direct interaction between MaHSF11 and MaDREB1, suggesting rather conserved genetic interaction between the two regulators (Supplemental Fig. S11). Recently, *MaCBF1* was reported to be a positive regulator of chlorophyll catabolism: It possesses strong transactivation activity and is localized to the nucleus (Xiao *et al*., 2023). It is also known to interact with MaNAC1 and involved in propylene-induced cold tolerance in banana (Shan *et al*., 2014). Overall, it may be concluded that MaHSF11 activates the expression of *MaDREB1* to fine tune flavonol synthesis. MaDREB1 may act, downstream of MaHSF11, to mediate drought, and cold stress response, as both are induced (Shan *et al*., 2014; Muthusamy *et al*., 2016). Prior to this study, several DREB proteins were reported to regulate different branches of flavonoid biosynthesis. CitERF32 and CitERF33 act as transcriptional inducers of *CitCHIL1* to promote flavone and flavanone biosynthesis in citrus (*Citrus reticulata cv. Suavissima*) (Zhao *et al*., 2021). In eggplant (*Solanum melongena*), SmCBF1, SmCBF2, and SmCBF3 interact with SmMYB113 to regulate *SmCHS* and *SmDFR* expression and enhance anthocyanin synthesis (Zhou *et al*., 2020). In our study, we discovered MaDREB1 to be a regulator of flavonol synthesis, modulating the expression of regulatory SG7 MYB genes. In eggplant as well, SmCBF1, SmCBF2, and SmCBF1 are induced by drought, salt, and ABA, showing that MaDREB1 might also be involved in drought stress signaling as a transcriptional regulator of stress-responsive genes such as flavonoid biosynthesis genes under drought and heat stress. Our study provides insights into a regulatory cascade that promotes flavonol biosynthesis in banana under different stress conditions, as shown in our proposed model (Fig. 9). HSF and DREB TFs have earlier been reported to modulate flavonol, and anthocyanin synthesis in few plant species under drought, cold and salt stress (Zhou *et al*., 2020; Bian *et al*., 2020; Wang *et al*., 2020). Previous studies have investigated the heat stress mediated modulation of flavonoids (Martinez *et al*., 2016; Zhou *et al*., 2021; Zhang *et al*., 2023; Yang *et al*., 2024). In our study, we explored the role of heat stress on flavonol synthesis, as HSFs are master regulators of heat stress signaling (Ohama *et al*., 2017). We found increased expression of *MaHSF11* and its downstream factors including MaDREB1, regulatory MYB and biosynthesis genes upon heat treatment. Further, increased accumulation of flavonol derivatives, particularly myricetin, upon heat stress verified the positive modulation of flavonol synthesis in banana by heat stress signalling, which corroborates with few previous studies (Martinez *et al*., 2016; Fragkostefanakis *et al*., 2016). Flavonol accumulation is promoted by high temperature in vegetative tissues and pollen in tomato (Martinez *et al*., 2016; Paupière *et al*., 2017). Banana genome encodes seven homologs of *MaF3’5’H* (Pandey *et al*., 2016). Among these few paralogs showed significantly increased transcript levels at early time points of 42□ heat treatment, while *MaF3’5’H4/6* transcript sharply increased at later time points. It might cumulatively contribute to increased myricetin accumulation upon heat treatment. In tomato, *HsfA2* mutant accumulates decreased transcripts of *CHI* which leads to reduced pollen viability under heat stress (Fragkostefanakis *et al*., 2016). In a recent study, a heat shock-induced B class HSF, MaHSF26 represses the expression of *MaRBOHs*, which leads to decreased ROS and MDA (Malondialdehyde) accumulation and alleviation of chilling injury in banana (Si *et al*., 2022). Our findings suggest rather consistent role of MaHSF11 under heat, drought, and salt stress mediated promotion of flavonol accumulation in banana (Fig. 2a, Fig. 8b, Wei *et al*., 2016).

**Figure 9.**
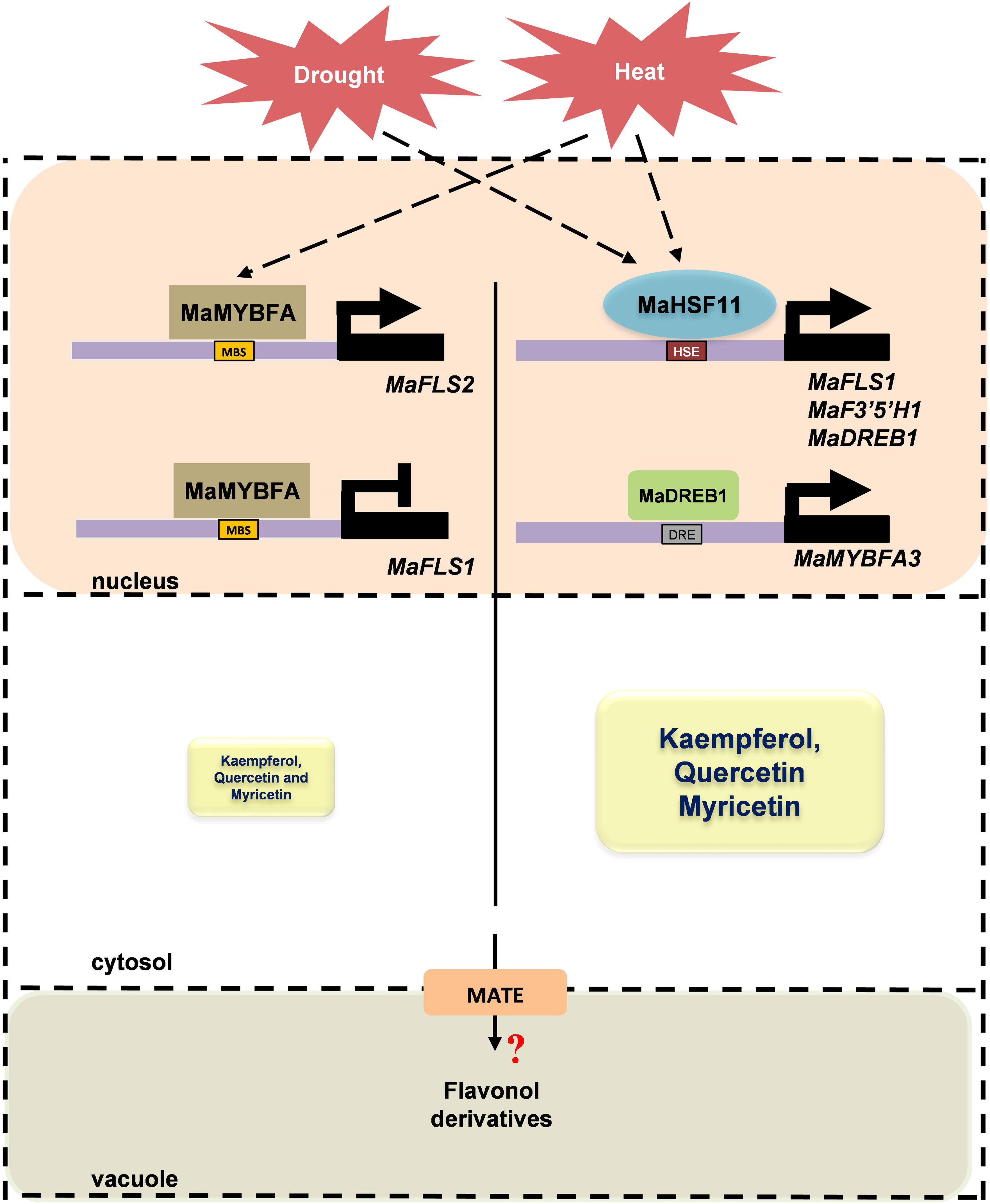
A working model showing the modulation of hydroxylated flavonol synthesis by the MaHSF11–MaDREB1–MaMYBFA regulatory module. The banana SG7 R2R3 MYB, MaMYBFA, targets *MaFLS2* to promote its expression, while strongly repressing the expression of *MaFLS1*, and modulate flavonol synthesis. Drought stress increases the transcript level of *MaHSF11*, enhancing the expression of *MaFLS1* and *MaF3*′*5*′*H1*. MaHSF11 also directly activates the expression of *MaDREB1*. MaDREB1 binds to the *MaMYBFA3* promoter to regulate its expression. This regulatory module enhances overall accumulation of flavonols under drought stress. MaMYBFA1, MaMYBFA2, and MaMYBFA3 differentially regulate the expression of *MaFLS1* and *MaFLS2*. MRE, MYB recognition element; DRE, dehydration response element; HSE, heat shock element. Heat stress modulates the expression of *MaHSF11, MaDREB1*, and *MaMYBFA1-3*, which subsequently leads to up-regulation of flavonol synthesis genes and increased accumulation of flavonols under heat stress in banana. Flavonols are likely transported into the vacuole by MATE transporters (multidrug and toxic compound extrusion), once the stress subsides.

The banana cultivar Grand Naine (Cavendish, AAA) is highly susceptible to different abiotic stresses as well as pathogen infection. Flavonols acting as ROS scavengers and pathogen defense molecules are synthesized in increased amounts under a variety of stresses. Here, we identified the HSF–DREB-MYB transcriptional regulatory module and characterized how it works independently or cooperatively to regulate flavonol synthesis in stress-responsive manner particularly, drought and heat conditions in banana. The TFs in this regulatory module can be used for effective engineering of stress resistance in crop plants like banana, as well as flavonol fortification of banana fruit. These candidate genes could be utilized for metabolic engineering of rutin and myricetin synthesis in banana fruit. Monocots are suggested to be monophyletic arising from a common ancestor (Remizowa and Sokoloff, 2023). It suggests conservation of regulatory mechanisms of stress-mediated flavonol synthesis in other monocot crops like, rice, maize, wheat. We have, in our study, explored a novel regulatory module of stress-mediated flavanol biosynthesis in banana. Which can also be applied or studied in other crops, particularly monocots.

## Materials and methods

### Plant material

Banana (cultivar subgroup Cavendish [AAA] variety ‘Grand Naine’) plants were grown in the fields of National Institute of Plant Genome Research (NIPGR), New Delhi, India. Various tissues, including young leaf, bract, pseudostem, root, peel, and pulp, of different fruit developmental stages were sampled during winter season, frozen in liquid nitrogen, and stored at −80□ until further use. *Nicotiana benthamiana* plants were grown in a Percival AR-41L3 growth cabinet at 100 µmol m^−2^ s^−1^ light intensity and 22°C under a 16-h-light/8-h-dark photoperiod.

### Identification of candidate flavonol-specific TFs from banana

To identify candidate SG7 R2R3-MYB TFs, which are known activators of flavonol biosynthesis, we retrieved sequences of previously characterized flavonol-specific TFs from various plants from NCBI (https://www.ncbi.nlm.nih.gov). These sequences were used to perform a BLASTp search against the banana genome sequence (DH-Pahang version 2) (https://banana-genome-hub.southgreen.fr). For phylogenetic studies, MEGA X (Kumar *et al*., 2018) was used to construct a maximum-likelihood phylogenetic tree with 1000 bootstrapping rounds. For multiple sequence alignment, MaMYBFA1–3 peptide sequences were aligned with a set of similar MYBs (Stracke *et al*., 2014) using MAFFT v.7.299b (Katoh and Standley, 2013). Different domains and motifs were identified manually (Naik *et al*., 2021).

### Cloning of flavonol-specific transcriptional regulators

The full-length CDSs (without the stop codons) of the MaMYB-encoding genes *MaMYBFA1* (*Ma02_g00290*), *MaMYBFA2* (*Ma05_g23640*), and *MaMYBFA3* (*Ma08_g10260*) were amplified using first-strand cDNA of banana fruit tissue and a set of primers containing Gateway attB sites that were designed based on information from the Banana Genome Hub (Supplemental Table S1). The resulting amplicons were recombined into the Gateway vector pDONR^TM^zeo (Invitrogen) and transformed into *Escherichia coli* TOP10 (Invitrogen) cells. Sanger sequencing of the resulting entry plasmids was performed by the NIPGR sequencing core facility (New Delhi, India).

### Subcellular localization and co-localization assays

CDSs of *MaMYBFA1–3, MaHSF11*, and *MaDREB1* were cloned into the pSITE-3CA binary vector (Chakrabarty *et al*., 2007) via LR recombination, producing yellow fluorescent protein (YFP) fusion proteins. Empty vectors were used as negative controls. The binary plasmids were transformed into *Agrobacterium tumefaciens* strain GV3101-pMP90 (Koncz & Schell,□1986). Equal volumes of *A. tumefaciens* cultures having the desired gene and the marker were mixed for agro-infiltration. *N. benthamiana* leaves were infiltrated on the abaxial side using a syringe, according to the protocol described by Walter *et al*. (2004). Infiltrated plants were kept in the dark at 25°C for 2 days prior to analysis. Leaf discs were then cut from the infiltrated sites and observed under an argon laser confocal laser-scanning microscope (TCS SP5; Leica Microsystems, Wetzlar, Germany) with YFP and RFP filters. YFP and RFP fluorescence was observed at 514-nm excitation/527-nm emission and 558-nm excitation/583-nm emission wavelengths, respectively. YFP/RFP gains (intensities %) for MaMYBFA1, MaMYBFA2, MaMYBFA3, MaDREB1 and MaHSF11 were 760 (39)/158 (39), 818 (39)/158 (39), 818 (30)/158 (39), 820 (45)/168(42) and 718 (42)/154 (39), respectively.

### RT-qPCR analysis

Total RNA was extracted from different banana organs according to a previously described protocol (Asif *et al*., 2006) and subsequently treated with RNase-free DNase (TURBO DNA-freeTM Kit, Invitrogen). Total RNA was reverse transcribed to generate first-strand cDNA using a RevertAid H Minus First Strand cDNA Synthesis Kit (Thermo Fisher) and oligo(dT) primers. RT-qPCR-based gene expression analyses were performed on a 7500 Fast Real-time PCR System (Applied Biosystems) using the 2x SYBR Green PCR Master mix (Applied Biosystems) and diluted cDNA corresponding to 10□ng total RNA in a final volume of 10 µL. *MaACTIN1* (GenBank AF246288) was used to normalize transcript abundance. Relative transcript levels were calculated using the cycle threshold (C_t_) 2^−ΔΔCT^ method (Livak & Schmittgen,□2001) and are presented as fold changes relative either to the tissue having the lowest expression or to the control condition, depending on the requirement. Three biological replicates or three technical replicates of a pool of 4-5 biological replicates were analyzed in each RT-qPCR analysis.

### Metabolite analysis

Targeted metabolite analysis was performed to quantify flavonols according to a protocol described previously (Pandey *et al*., 2016; Naik *et al*., 2021; Rajput *et al*., 2022a, b). Plant metabolites were extracted from shredded plant material in 80% methanol at room temperature. Finely ground tissues were dissolved in 80% methanol, kept at 70□for 15 minutes, and subjected to centrifugation for 10 minutes at 12,000 rpm (18514 g). After centrifugation, supernatants were mixed with three volumes of 1N HCl for acid hydrolysis and incubated at 94□for 2 h. After incubation, an equal volume of ethyl acetate was added, and the upper phase was removed for further purification of the desired metabolites. The ethyl acetate was evaporated at 37□ under lower pressure (90–100 mbar) on a Buchi rotavapor R-300 (BUCHI, Switzerland), and samples were finally dissolved in 80% methanol. Liquid chromatography–mass spectrometry (LC-MS) analysis was performed using an ultra-high-performance liquid chromatography (UHPLC) system (Exion LC Sciex, Framingham) coupled to a triple quadrupole system (QTRAP6500+; ABSciex) with electrospray ionization. The voltage was set to 5500□V for positive ionization. Detailed UHPLC and LC-MS parameters have been described by Naik *et al*. (2021). Identification and quantitative analyses were done via Analyst software (version 1.5.2).

### Cloning of reporter constructs and identification of putative *cis* elements in promoters

Total genomic DNA was extracted from banana leaves using the DNeasy Plant Mini Kit (Qiagen). Sequences of the putative target gene promoter fragments were retrieved from the banana genome (https://banana-genome-hub.southgreen.fr) (D’Hont *et al*., 2016) and analyzed for the presence of *cis*-regulatory motifs via New PLACE software (https://www.dna.affrc.go.jp) supplemented by manual curation. Promoter fragments were amplified from banana genomic DNA using designed primer sequences and Phusion™ High-Fidelity DNA Polymerase (Thermo Scientific). The amplified fragments were cloned into the pENTR™/D-TOPO™ vector (ThermoFisher) and sequenced to confirm their integrity. The entry clones of promoters (*proMaFLS1–2508*, *proMaF3*□*5*′*H1–1335*, *proMaDREB1–998*, and *proMaMYBFA3–2508*) were recombined into the vector p635nRRF containing *pro35S:REN* (Kumar *et al*., 2018) to make the reporter constructs.

### Cloning of cDNA from MaHSF and MaDREB transcriptional regulators

Full-length CDSs of selected homologs of MaHSF and MaDREB were amplified by first-strand cDNA synthesis of banana leaves using Phusion™ High-Fidelity DNA Polymerase (Thermo Scientific) and a set of primers containing Gateway attB sites designed on the basis of information from the Banana Genome Hub (Supplemental Table S1). The resulting amplicons were cloned into the pDONR^TM^ zeo vector (Invitrogen) and transformed into *E. coli* TOP10 cells, and integrity was confirmed by Sanger sequencing (Sanger *et al*., 1977).

### Dual-luciferase reporter assay

To test the transactivation potential of the transcriptional regulators, MaHSF11, and MaDREB1 CDSs were recombined into the destination vector pBTdest (Baudry *et al*., 2004) to generate CaMV *pro35S*-driven effector constructs. For dual-luciferase reporter assays, reporter and effector constructs were transformed into *A. tumefaciens* strain GV3101 (pMP90) (GoldBio) and subsequently infiltrated using a syringe into the abaxial side of *N. benthamiana* leaves according to a previously described protocol (Walter *et al*., 2004). LUC and REN activities were measured 48□h later in extracts from infiltrated leaf discs using the Dual-Luciferase Reporter Assay System (Promega) according to the manufacturer’s protocol. Luminescence in the samples was quantified using the POLARstar Omega multimode plate reader (BMG Labtech), and LUC activity was normalized to REN activity. Values are given as means ± SD of the LUC/REN ratio of 4 independent biological replicates.

### Y1H assays

The Matchmaker Gold Yeast 1-hybrid system (Clontech) with the yeast one-hybrid (Y1H) gold yeast strain was used. Promoter fragments were cloned into the pABAi vector (Takara Bio), and the pABAi plasmids carrying the promoter fragments were linearized and transformed into the Y1H Gold strain according to the Matchmaker protocol, whereby they integrated into the yeast genome by homologous recombination. Effector plasmids and empty pGADT7-GW vector (Gal4-AD fusion, Ura selection) controls were transformed into the Y1H strains previously transformed with AurR-reporter constructs (*proMaFLS1*-AurR, *proMaF3*′*5*′*H1*-AurR, *proMaDREB1*-AurR, and *proMaMYBFA3*-AurR). To test for interaction, the yeast colonies containing effector and reporter constructs were spotted onto medium containing AbA per the manufacturer’s protocol. Growth of different Ura^−^ yeast colonies was checked using different concentrations of AbA to determine the minimum inhibitory concentration (MIC) values and to check for possible interactions. The MIC values for *proMaFLS1*, *proMaF3*′*5*′*H1*, *proMaDREB1*, and *proMaMYBFA3* were 100□ng□mL^−1^ AbA, 1000□ng□mL^−1^ AbA, 100□ng□mL^−1^, and 900□ng□mL^−1^, respectively.

### ChIP-qPCR assay

The ChIP-qPCR assay was performed using the ChIP Kit – Plants kit (Abcam) according to the manufacturer’s instructions (Shi *et al*., 2019). Promoter fragments containing the HSE *cis* motif in p635nRRF that was used for dual-luciferase assays were co-infiltrated with YFP-MaHSF11 fusion proteins into *N. benthamiana* leaves. Three days after infiltration, equally sized leaf discs were used for ChIP. Next, 2 µL anti-GFP antibody (Catalog #ab290, Abcam) was used for chromatin precipitation in two biological replicates, and 2 µL of anti-IgG antibody (Catalog #ab48386, Abcam) was used as a negative control. qPCR was performed with 5 µL 2x SYBR Green PCR Master mix (Applied Biosystems), 1 µL ChIP DNA, and 0.5 µL each forward and reverse primer, with 40 cycles of amplification as described above. The relative fold enrichment of the target promoter fragments was calculated using the 2^−(–ΔCT)^ method (Livak & Schmittgen, 2001). The primers used in ChIP-qPCR are listed in Supplemental Table S1.

### Electrophoretic mobility-shift assay

For the EMSA, the fusion construct MaHSF11-GST or MdDREB1-GST with GST-tag were obtained by cloning the CDS of MaHSF11 or MaDREB1 into pGEX-4T-2 vector, and then individually introduced into *Escherichia coli* cells BL21 (DE3/Codon+) for protein production. The culture solution was added with IPTG (final concentration 0.5 mM) at OD of 0.6 and kept on culturing at 28 □, for 12 h followed by harvesting. GST-tagged recombinant protein was purified using Glutathione Resin (catalog no. 786-310, G-Biosciences). The probes containing intact and mutated cis-motifs were labelled with biotin using Pierce™ Biotin 3’ End DNA Labeling Kit (catalog no. 89818, Thermo Fisher Scientific, USA). The EMSA was carried out using LightShift Chemiluminescent EMSA Kit (catalog no. 20148, Thermo Fisher Scientific, USA) according to the protocol described in the kit manual. The primers and probe sequences for EMSA are shown in Supplementary Table S1.

### Transient expression in banana fruit

For transient overexpression studies, the CDSs of *MaMYBFA1*, *MaMYBFA2*, *MaMYBFA3*, *MaHSF11*, and *MaDREB1* were cloned into pANIC6b (Mann *et. al*., 2012), a Gateway-compatible destination vector suitable for overexpression of genes of interest using *A. tumefaciens*-mediated transformation, particularly useful for monocot species. pANIC6b encodes hygromycin resistance under the control of a constitutive rice (*Oryza sativa*) *actin1* promoter (*proOsACT1*) (LOC4338914), a constitutive *Panicum virgatum ubiquitin1+3* promoter (*proPvUBI*)-driven GUSPlus visual marker, and a Gateway cassette driven by a *Zea mays ubiquitin1* promoter and intron for overexpression of target genes. The resulting overexpression plasmid and the empty vector control were transformed into *A. tumefaciens* strain EHA105 (Mehra *et al*., 2019). Resulting *A. tumefaciens* suspensions were then separately introduced into discs of immature green banana fruits by vacuum infiltration as previously described (Rajput *et al*., 2022a). After 3 days of co-cultivation on MS media (MS basal salt mixture, HiMedia), transformation of banana fruit disc was confirmed by GUS staining. Tissues were harvested and frozen in liquid nitrogen for subsequent expression (RT-qPCR) and metabolite (LC-MS) analyses.

For gene silencing through RNA interference, the 250-350 bp CDS fragments of *MaHSF11* and *MaDREB1* were cloned into pANIC8b vector, a Gateway-compatible destination vector suitable for silencing of target genes in monocot species (Mehra *et al*., 2022). The transformation into *A. tumefaciens* strain EHA105 and immature banana fruits were done as described previously for overexpression. After GUS staining, tissues were harvested for expression (RT-qPCR) and metabolite analysis (LC-MS).

### Heat stress treatments

We used plant growth chamber (Model: AR-41L3, Percival) to perform heat stress treatment at 250 µmol m^−2^ s^−1^ light intensity under a 16-h-light/8-h-dark photoperiod and 60% relative humidity. Three months old *in vitro* grown banana plants (cultivar subgroup Cavendish [AAA] variety ‘Grand Naine’) of equal size were kept at 26°C for control and 42°C for heat treatment. Leaf samples were harvested at 0, 6, 12, 24, 48, and 72 hours (h) post heat stress treatment followed by freezing in liquid nitrogen. Expression (RT-qPCR) and metabolite analysis (LC-MS) were carried out as described earlier.

### Y2H assay

For Y2H assays, full lengthen CDS of MaHSF11 and MaDREB1 were cloned into pGADT7 (Clontech Laboratories Inc., San Jose, CA,□USA) and pGBKT7 vector, respectively. The resulting AD and BD constructs were co-transformed into Y2HGold (Clontech) and grown at 30°C on synthetic defined (SD) medium devoid of Trp and Leu. For interaction assay, the colonies having both the bait and prey clones were spotted onto medium lacking Trp, Leu, Ade, and His (QDO). p53/large T antigen interaction (Pipas & Levine,□2001) was used as the positive control and empty pGADT7g/pGBKT7g vectors with opposite and appropriate constructs were used as a negative control.

### Statistical tests

One-way and two-way analysis of variance (ANOVA) with Tukey’s multiple comparison were performed for statistical analysis. Unpaired Student’s *t*-tests were used for the specific comparisons. Asterisks indicate the level of significance: *\*P*□≤□0.05; *\*\*P*□≤□0.01; *\*\*\*P*□≤□0.001, *\*\*\*\*P* ≤ 0.0001; ns, not significant.

## Data availability

All data supporting the findings of this study are available within the paper and within the supplemental data published online.

## Supplemental data

**Figure S1.** Identification of candidate R2R3-MYB flavonol regulators from the banana genome and their subcellular localization.

**Figure S2.** Tissue-specific expression of *MaMYBFA1-FA3, MaFLS1-2* genes and flavonol content in different tissues of banana.

**Figure S3.** Relative expression of structural flavonol biosynthesis gene transcripts in banana fruit discs transiently overexpressing *MaMYBFA1-FA3*.

**Figure S4.** Relative expression of structural and regulatory flavonol biosynthesis genes in *MaHSF11* overexpressing banana fruit discs.

**Figure S5.** Relative expression of structural and regulatory flavonol biosynthesis genes in *MaHSF11*-silenced banana fruit discs.

**Figure S6.** Phylogenetic analysis of MaDREB1 with previously characterized DREB proteins from other plant species, its sub-cellular localization and tissue specific expression pattern in different tissues of banana.

**Figure S7.** Relative expression of structural flavonol biosynthesis genes in overexpressing *MaDREB1* overexpressing banana fruit discs.

**Figure S8.** Relative expression of structural and regulatory flavonol biosynthesis genes in *MaDREB1*-silenced banana fruit discs.

**Figure S9.** Relative expression analysis of flavonol biosynthesis genes in control (26 L) and heat stress (42 L).

**Figure S10.** Relative expression analysis of *MaHSF11* in overexpressing *MaDREB1*– and *MaDREB1*-silenced banana fruit discs.

**Figure S11.** Interaction between MaHSF11 and MaDREB1 in yeast.

## Acknowledgements

This work was supported by a core grant from the National Institute of Plant Genome Research and a Department of Science and Technology-SERB for Core research grant (CRG/2022/001178) to AP. JN, RR and SS acknowledge the Council of Scientific and Industrial Research, Government of India, for Senior Research Fellowships. The authors thank the DBT-eLibrary Consortium (DeLCON) for providing access to e-resources. We also acknowledge the Metabolome facility at NIPGR for phytochemical analysis.

## Declarations

The authors declare no conflicts of interest.

## Supplemental Figures

**Supplemental Figure S1.** Identification of candidate R2R3-MYB flavonol regulators from the banana genome and their subcellular localization. (a) Various R2R3-MYB proteins were classified into three major groups as flavonol-, anthocyanin-, or proanthocyanidin biosynthesis–specific regulators using landmark flavonoid-specific R2R3-MYBs from different plant species. Numbers on each branch show branch length, and solid dark blue circles represent bootstrap values. (b) Amino acid sequence alignment of candidate SG7 R2R3-MYB transcription factors (MaMYBFA1, MaMYBFA2, and MaMYBFA3, where FA stands for flavonol activator) and selected flavonol regulators from other plant species. The R2 and R3 domains are highlighted in grey. The SG7-1 and SG7-2 motifs, found in subgroup 7 R2R3-MYBs, are highlighted in dark green and yellow, respectively. Accession numbers and two-letter species abbreviations are provided in Supplemental File S2. (c) Subcellular localization of MaMYBFAs. MaMYBFA-YFP fusion proteins were expressed together with the nuclear marker NLS-RFP in *N. benthamiana* leaf epidermal cells. Confocal imaging shows co-localization of YFP and RFP fluorescence, indicating nuclear localization of the three MaMYBFA proteins. Empty vector pSITE-3CA was also taken as negative control, which shows free YFP. Representative images are shown with scale bar of 50 µm.

**Supplemental Figure S2.** Tissue-specific expression of *MaMYBFA1–3, MaFLS1-2* genes and flavonol content in different tissues of banana. (a) Relative expression of *MaMYBFA1–3* genes in different tissues of banana as determined by RT-qPCR. (b) Relative expression of *MaFLS1* and *MaFLS2* genes in different tissues of banana. (c) Flavonol aglycones i.e., kaempferol, quercetin and myricetin content in µg per 100g dry weight (DW) in different tissues of banana. Tissues from three independent biological replicates were used for expression and metabolite (LC-MS) analysis. YL, young leaf; BRT, bract; P.STEM, pseudostem; UR.PL, unripe peel; UR.PP, unripe pulp; R.PL, ripe peel; R.PP, ripe pulp. Values shown are means ± SD of three biological replicates.

**Supplemental Figure S3.** Relative expression of structural flavonol biosynthesis gene transcripts in banana fruit discs transiently overexpressing *MaMYBFA1-FA3*. RT-qPCR-based transcript profiling of structural flavonol biosynthesis genes *MaCHS1–6*, *MaCHI1–2*, *MaF3H1–2*, *MaF3*′*5*′*H1–7*, and *MaF3*′*H* in *MaMYBFA1–3* OE banana fruit discs. EV, empty vector control. Values shown are means ± SD of three technical replicates of pool of 4-5 independent biological samples. One-way ANOVA was used for statistical analysis with Tukey’s multiple comparison. \**P*□≤□0.05; \*\**P*□≤□0.01; \*\*\**P*□≤□0.001; \*\*\*\**P* ≤ 0.0001; ns, not significant.

**Supplemental Figure S4.** Relative expression of structural and regulatory flavonol biosynthesis genes in *MaHSF11* overexpressing banana fruit discs. RT-qPCR-based expression analysis of (a) *MaCHS1–6*, (b) *MaCHI1–2*, (c) *MaF3H1–2*, (d) *MaF3*′*H*, and (e) *MaMYBFA1–3* in *MaHSF11* overexpressing banana fruit discs. Values shown are means ± SD of three technical replicates of a pool of 4-5 independent biological samples. EV, empty vector control. Two-way ANOVA was used with Tukey’s multiple comparison. *P*□≤□0.05; *\*\*P*□≤□0.01; ns, not significant. N.D., not determined.

**Supplemental Figure S5.** Relative expression of structural and regulatory flavonol biosynthesis genes in *MaHSF11*-silenced banana fruit discs. RT-qPCR-based expression analysis of (a) *MaCHS1–6*, (b) *MaCHI1–2*, (c) *MaF3H1–2*, (d) *MaF3*′*H*, and (e) *MaMYBFA1–3* in EV and *proZmUBI:MaHSF11-RNAi* banana fruit discs. Values shown are means ± SD of three technical replicates of a pool of 4-5 independent biological samples. EV, empty vector control. Two-way ANOVA was used with Tukey’s multiple comparison. \**P*□≤□0.05; *\*\*P*□≤□0.01; *\*\*\*\*P* ≤ *0.0001;* ns, not significant. N.D., not determined.

**Supplemental Figure S6.** Phylogenetic analysis of MaDREB1 with previously characterized DREB proteins from other plant species, its sub-cellular localization and tissue specific expression pattern in different tissues of banana. (a) The phylogenetic tree was constructed in MEGA-X32 software using the maximum-likelihood method with 1000 bootstrap values and visualized with iTOL v.6.4 (http://itol.embl.de/). Accessions used were AtDREB1B (AAV80413.1), AtDREB1C (NP_567719.1), AtDREB1C (AAV80414.1), GmDREB3 (AAZ03388.1), GmDREB2 (NP_001237254.2), GmDREB2A2 (NP_001240942.3), OsDREB1 (AAL40870.1), OsDREB1B (AAN85707.1), OsDREB3 (ADO79620.1), SlDREB3 (NP_001234707.1), HvDREB2 (AFH75401.1), and ZmDREB (NP_001105081.1). At, *Arabidopsis thaliana*; Gm, *Glycine max*; Os, *Oryza sativa*; Sl, *Solanum lycopersicum*; Hv, *Hordeum vulgare*; Zm, *Zea mays*). (b) Subcellular localization of MaDREB1 in *N. benthamiana* leaves. The YFP-MaDREB1 fusion protein localizes to the nucleus and coincides with the signal from the nuclear localization signal-red fluorescent protein (NLS-RFP) fusion in *N. benthamiana* leaves as observed in confocal microscopy. Empty vector pSITE-3CA as used as negative control to visualize free YFP. Scale bar, 50Lμm. (c) Relative expression of *MaDREB1* gene in different tissues of banana as determined by RT-qPCR. Values shown are means ± SD of three biological replicates.

**Supplemental Figure S7.** Relative expression of structural flavonol biosynthesis genes in *MaDREB1* overexpressing banana fruit discs. RT-qPCR-based expression analysis of (a) *MaCHS1–6*, (b) *MaCHI1–2*, (c) *MaF3H1–2*, (d) *MaF3’H* and (e) *MaF3*′*5*′*H1–7* in *MaDREB1* overexpressing banana fruit discs. Values shown are means ± SD of three technical replicates of a pool of 4-5 independent biological pools. Two-way ANOVA was used with Tukey’s multiple comparison*. *P*□≤□0.05; *\*\*P*□≤□0.01; *\*\*\*P*□≤□0.001; ns, not significant.

**Supplemental Figure S8.** Relative expression of structural flavonol biosynthesis genes in *MaDREB1*-silenced banana fruit discs. RT-qPCR-based expression analysis of (a) *MaCHS1–6*, (b) *MaCHI1–2*, (c) *MaF3H1–2*, (d) *MaF3*′*H* and (e) *MaF3*′*5*′*H1–7* in *MaDREB1*-silenced banana fruit discs. Values shown are means ± SD of three technical replicates of a pool of 4-5 independent biological pools. Two-way ANOVA was used with Tukey’s multiple comparison. *PL≤L0.05; **PL≤L0.01; ****PL≤L0.0001; ns, not significant.

**Supplemental Figure S9.** Relative expression analysis of other flavonol biosynthesis genes in control (26□) and heat stress (42□) treated banana seedlings. RT-qPCR based relative expression level of *MaCHS1-6, MaCHI1-2, MaF3H1-2, MaF3’H*, and *MaF3’5’H1-7* at different time points (0h, 6h, 12h, 24h, 48h, 72h) of heat stress (42 □) treatment in banana seedlings, and control (26 L) seedlings. Values shown are means ± SD of three biological replicates.

**Supplemental Figure S10.** Relative expression analysis of *MaHSF11* in *MaDREB1* overexpressing *and MaDREB1*-silenced banana fruit discs. Values shown are means ± SD of three technical replicates of a pool of 4-5 independent biological samples. Unpaired Student’s *t*-tests were used with Tukey’s multiple comparison for statistical analysis. *\*\*\*P*□≤□0.001, ns. Not significant.

**Supplemental Figure S11.** Interaction between MaHSF11 and MaDREB1 in yeast. Yeast-two-hybrid assays showing no direct interaction of MaHSF11 with MaDREB1 in yeast. AD, GAL4 activation domain; BD, GAL4 DNA-binding domain; –LT medium, synthetic defined (SD)−Leu−Trp medium; –AHLT, SD–Leu–Trp–Ade–His medium.

**Supplemental File S1.** Promoter sequences with highlighted *cis* motifs used in this study.

**Supplemental File S2.** Accession numbers and two-letter species abbreviations used in phylogenetic tree figures.

**Supplemental Table S1.** List of primers used in this study.

