## Supplementary Figure for "The HSF-DREB-MYB transcriptional regulatory module regulates flavonol biosynthesis and flavonoid B-ring hydroxylation in banana (*Musa acuminata*)"

### Slide 1
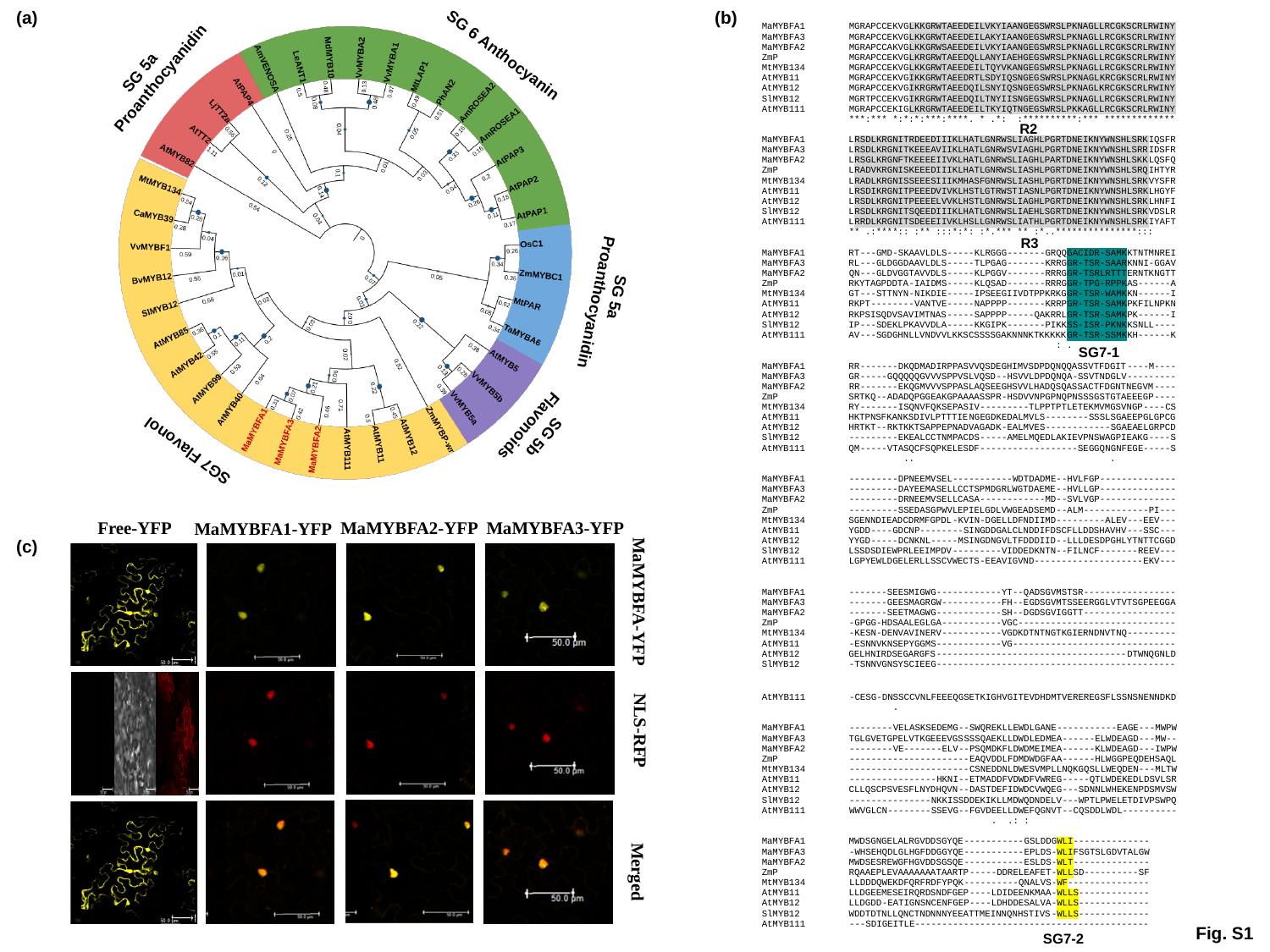

(a)
(b)
 SG 5a Proanthocyanidin
SG 6 Anthocyanin
R2
R3
 SG 5a Proanthocyanidin
SG7-1
 SG7 Flavonol
 SG 5b
 Flavonoids
Free-YFP
MaMYBFA2-YFP
MaMYBFA3-YFP
MaMYBFA1-YFP
(c)
MaMYBFA-YFP
NLS-RFP
Merged
Fig. S1
SG7-2

### Slide 2
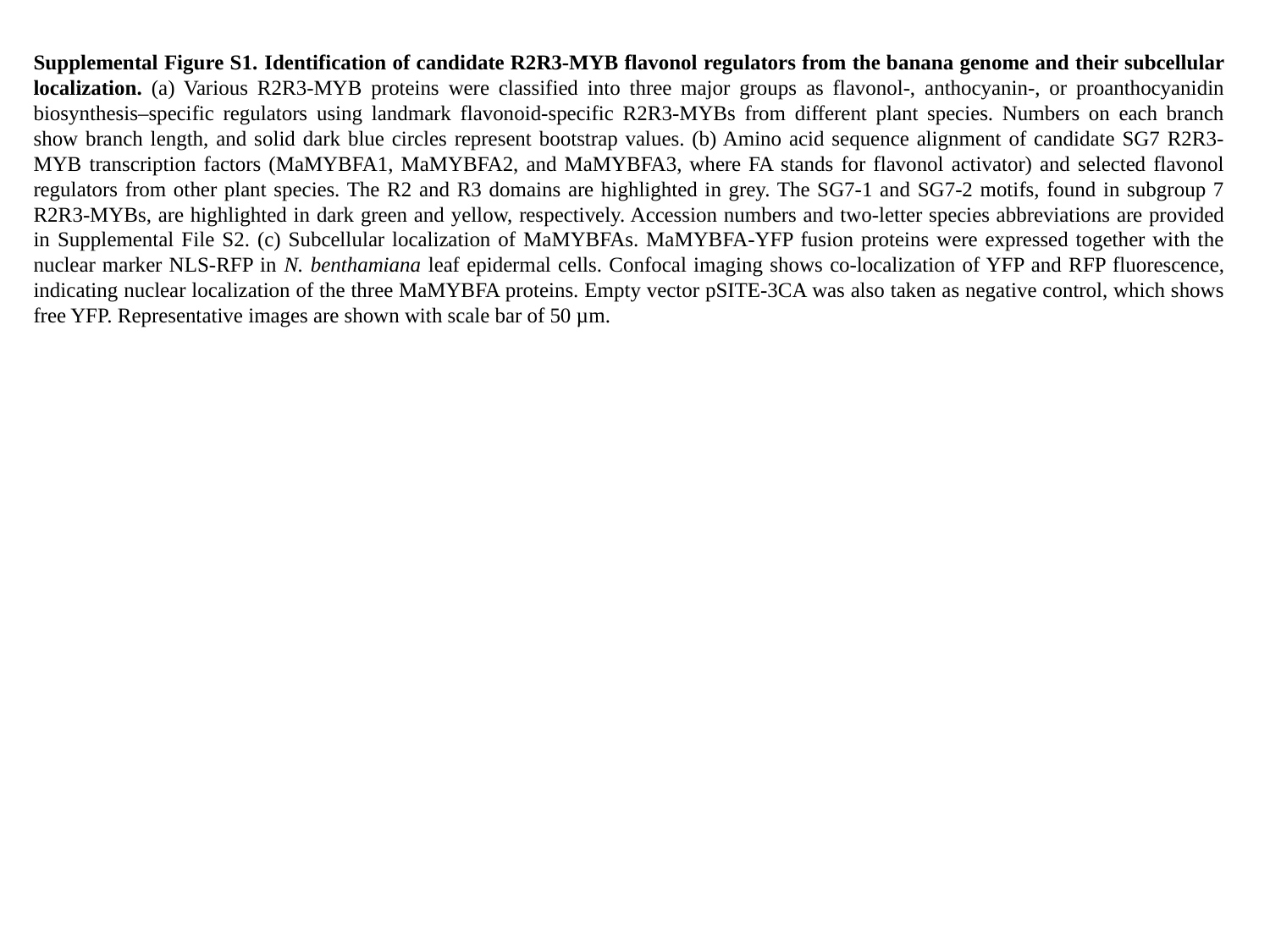

Supplemental Figure S1. Identification of candidate R2R3-MYB flavonol regulators from the banana genome and their subcellular localization. (a) Various R2R3-MYB proteins were classified into three major groups as flavonol-, anthocyanin-, or proanthocyanidin biosynthesis–specific regulators using landmark flavonoid-specific R2R3-MYBs from different plant species. Numbers on each branch show branch length, and solid dark blue circles represent bootstrap values. (b) Amino acid sequence alignment of candidate SG7 R2R3-MYB transcription factors (MaMYBFA1, MaMYBFA2, and MaMYBFA3, where FA stands for flavonol activator) and selected flavonol regulators from other plant species. The R2 and R3 domains are highlighted in grey. The SG7-1 and SG7-2 motifs, found in subgroup 7 R2R3-MYBs, are highlighted in dark green and yellow, respectively. Accession numbers and two-letter species abbreviations are provided in Supplemental File S2. (c) Subcellular localization of MaMYBFAs. MaMYBFA-YFP fusion proteins were expressed together with the nuclear marker NLS-RFP in N. benthamiana leaf epidermal cells. Confocal imaging shows co-localization of YFP and RFP fluorescence, indicating nuclear localization of the three MaMYBFA proteins. Empty vector pSITE-3CA was also taken as negative control, which shows free YFP. Representative images are shown with scale bar of 50 µm.

### Slide 3
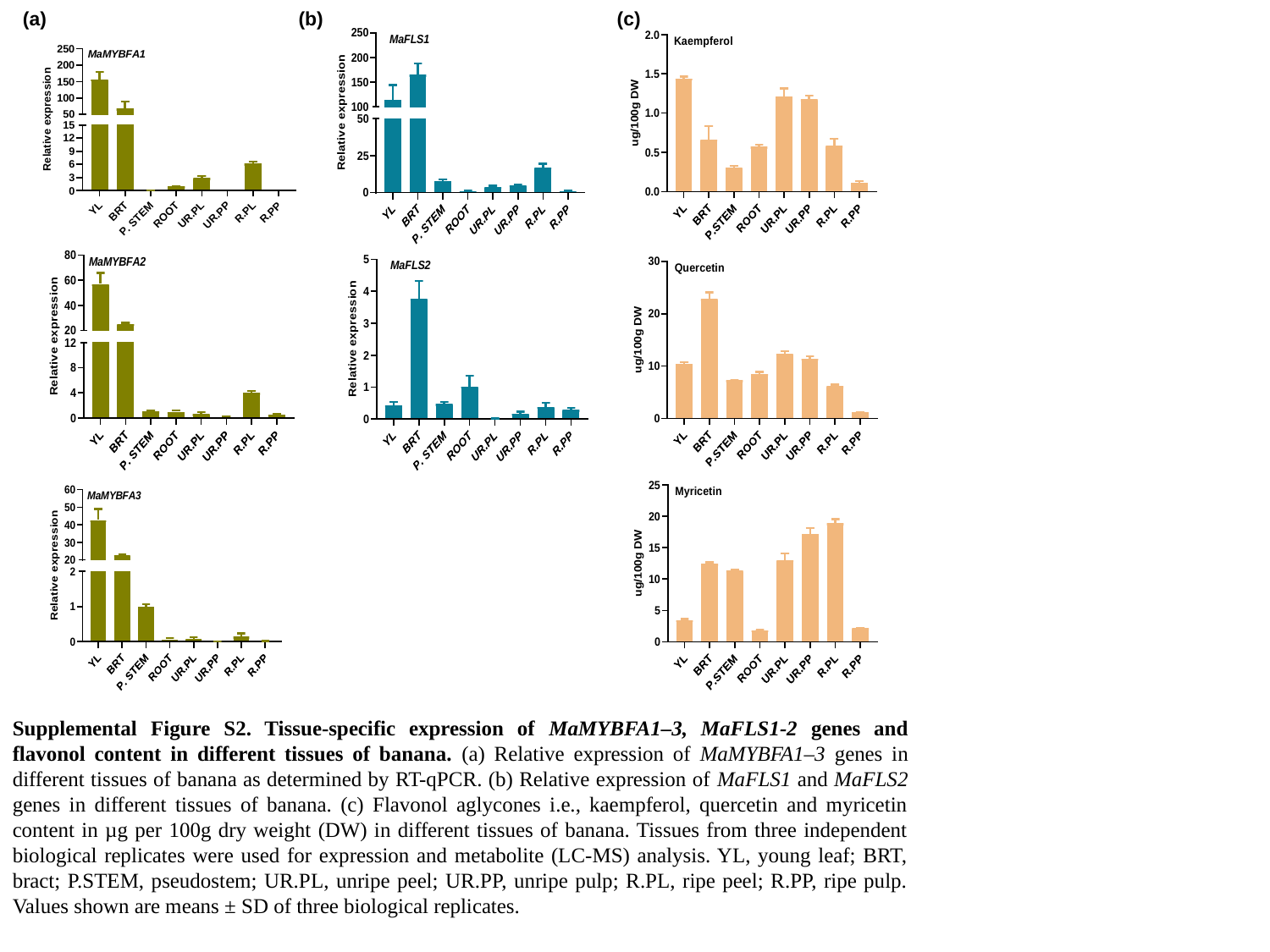

(a)
(b)
(c)
Supplemental Figure S2. Tissue-specific expression of MaMYBFA1–3, MaFLS1-2 genes and flavonol content in different tissues of banana. (a) Relative expression of MaMYBFA1–3 genes in different tissues of banana as determined by RT-qPCR. (b) Relative expression of MaFLS1 and MaFLS2 genes in different tissues of banana. (c) Flavonol aglycones i.e., kaempferol, quercetin and myricetin content in µg per 100g dry weight (DW) in different tissues of banana. Tissues from three independent biological replicates were used for expression and metabolite (LC-MS) analysis. YL, young leaf; BRT, bract; P.STEM, pseudostem; UR.PL, unripe peel; UR.PP, unripe pulp; R.PL, ripe peel; R.PP, ripe pulp. Values shown are means ± SD of three biological replicates.

### Slide 4
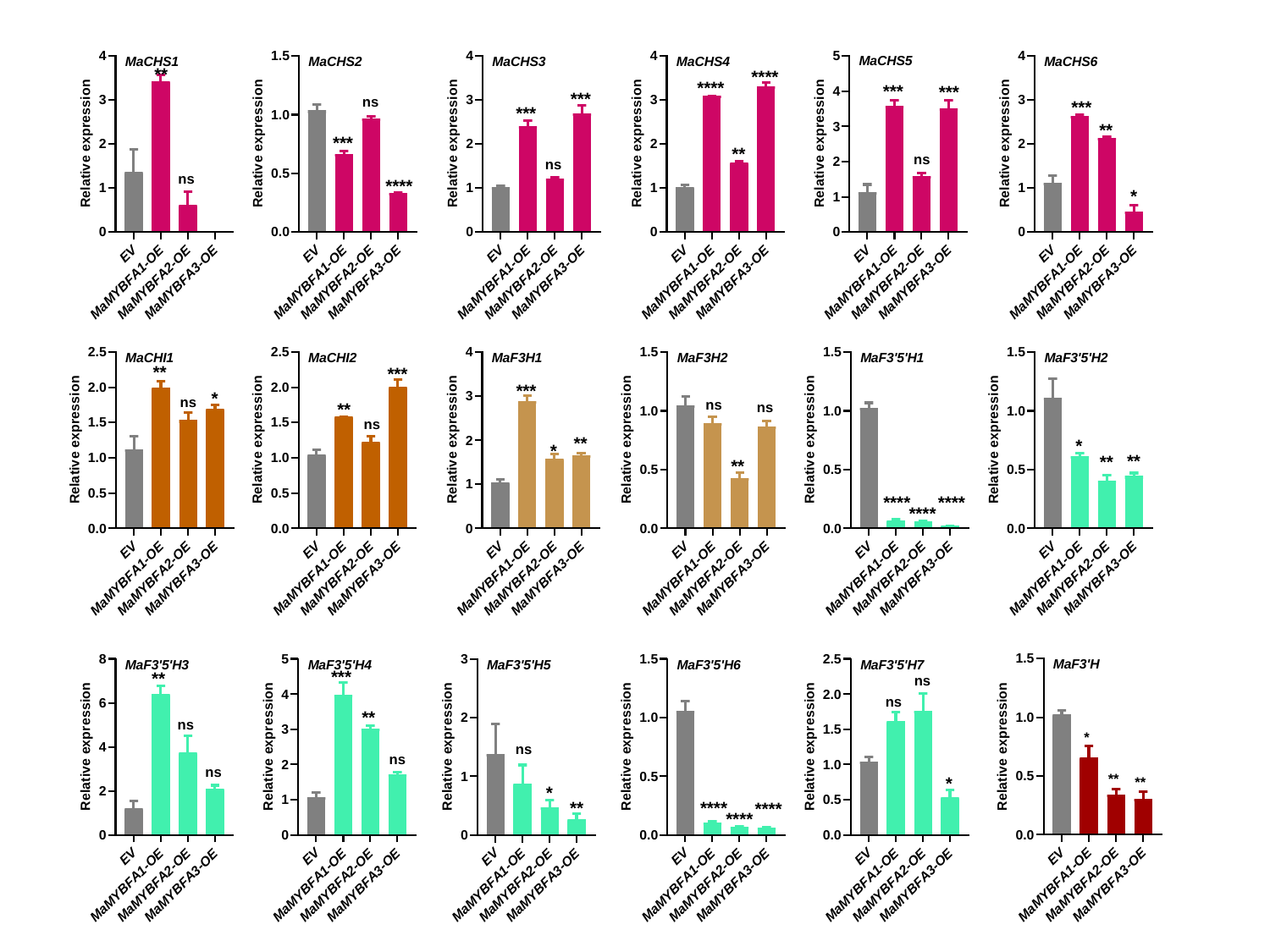

**
****
****
***
***
***
ns
***
***
**
***
**
ns
ns
ns
****
*
**
***
***
*
ns
ns
ns
**
ns
**
*
*
**
**
**
****
****
****
***
**
ns
ns
**
ns
*
ns
ns
ns
**
*
**
*
**
****
****
****

### Slide 5
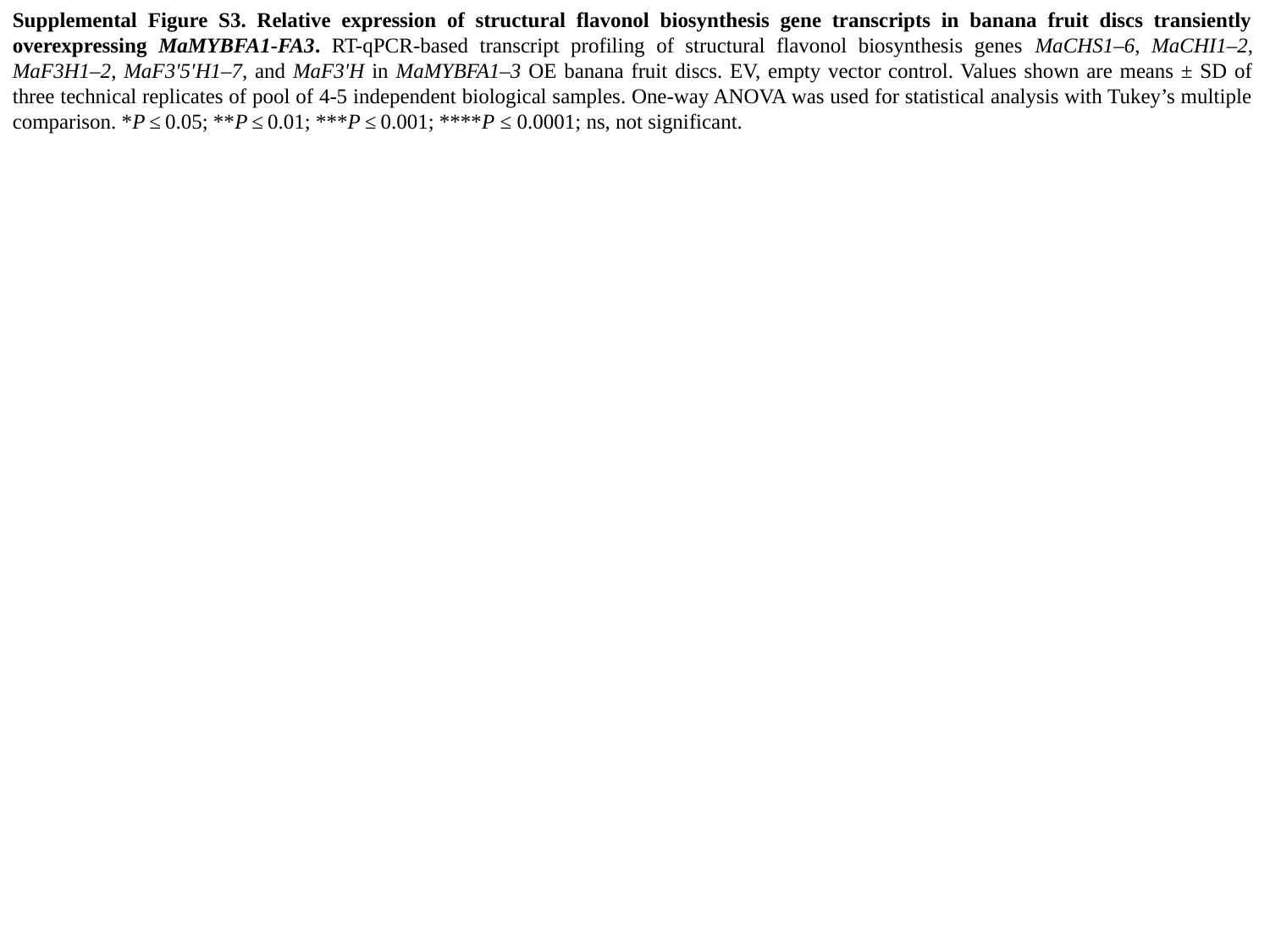

Supplemental Figure S3. Relative expression of structural flavonol biosynthesis gene transcripts in banana fruit discs transiently overexpressing MaMYBFA1-FA3. RT-qPCR-based transcript profiling of structural flavonol biosynthesis genes MaCHS1–6, MaCHI1–2, MaF3H1–2, MaF3′5′H1–7, and MaF3′H in MaMYBFA1–3 OE banana fruit discs. EV, empty vector control. Values shown are means ± SD of three technical replicates of pool of 4-5 independent biological samples. One-way ANOVA was used for statistical analysis with Tukey’s multiple comparison. *P ≤ 0.05; **P ≤ 0.01; ***P ≤ 0.001; ****P ≤ 0.0001; ns, not significant.

### Slide 6
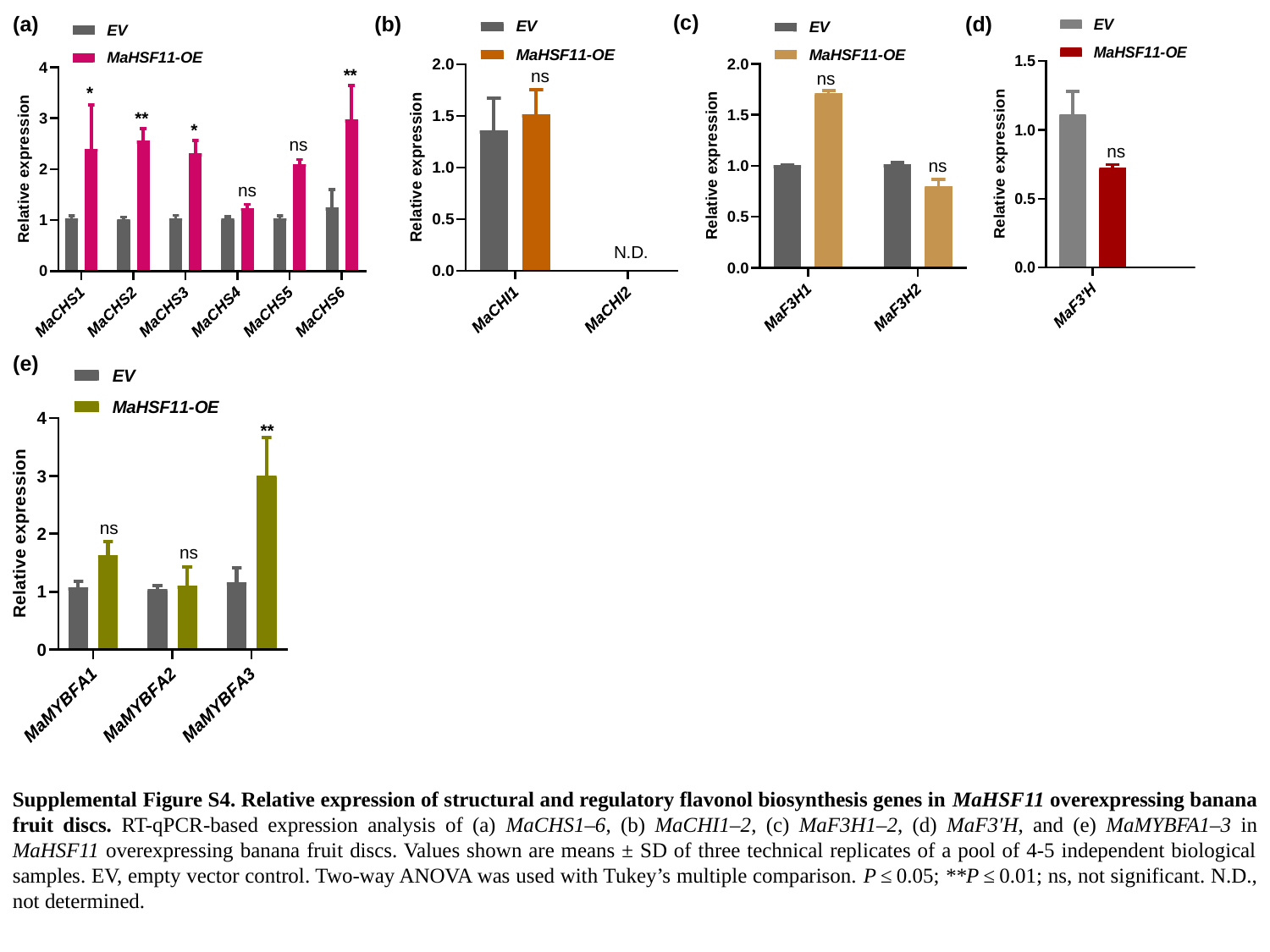

(c)
(b)
(a)
(d)
**
ns
ns
*
**
*
ns
ns
ns
ns
(e)
**
ns
ns
Supplemental Figure S4. Relative expression of structural and regulatory flavonol biosynthesis genes in MaHSF11 overexpressing banana fruit discs. RT-qPCR-based expression analysis of (a) MaCHS1–6, (b) MaCHI1–2, (c) MaF3H1–2, (d) MaF3′H, and (e) MaMYBFA1–3 in MaHSF11 overexpressing banana fruit discs. Values shown are means ± SD of three technical replicates of a pool of 4-5 independent biological samples. EV, empty vector control. Two-way ANOVA was used with Tukey’s multiple comparison. P ≤ 0.05; **P ≤ 0.01; ns, not significant. N.D., not determined.

### Slide 7
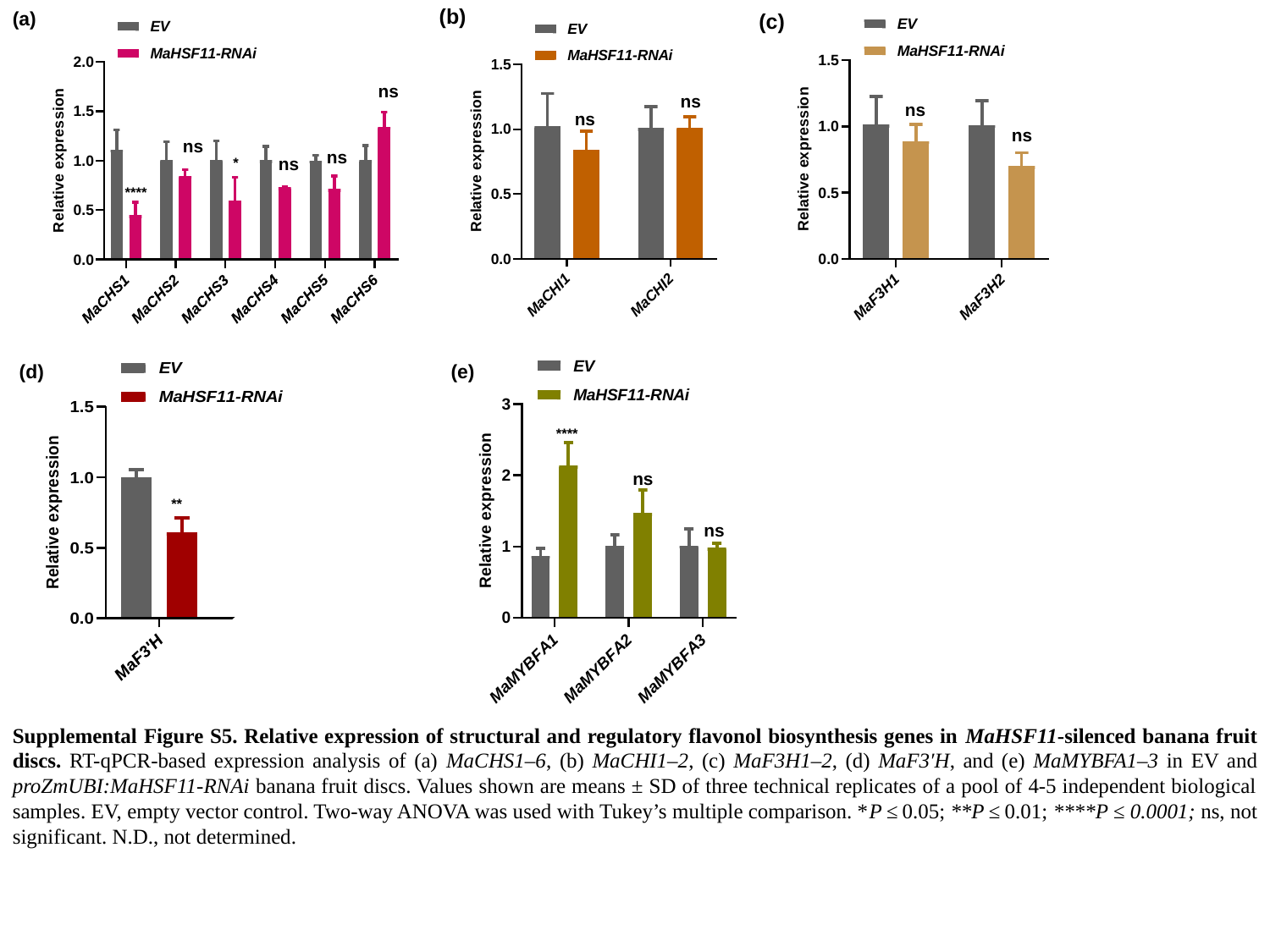

(a)
(b)
(c)
ns
ns
ns
ns
ns
ns
ns
ns
*
****
(d)
(e)
****
ns
**
ns
Supplemental Figure S5. Relative expression of structural and regulatory flavonol biosynthesis genes in MaHSF11-silenced banana fruit discs. RT-qPCR-based expression analysis of (a) MaCHS1–6, (b) MaCHI1–2, (c) MaF3H1–2, (d) MaF3′H, and (e) MaMYBFA1–3 in EV and proZmUBI:MaHSF11-RNAi banana fruit discs. Values shown are means ± SD of three technical replicates of a pool of 4-5 independent biological samples. EV, empty vector control. Two-way ANOVA was used with Tukey’s multiple comparison. *P ≤ 0.05; **P ≤ 0.01; ****P ≤ 0.0001; ns, not significant. N.D., not determined.

### Slide 8
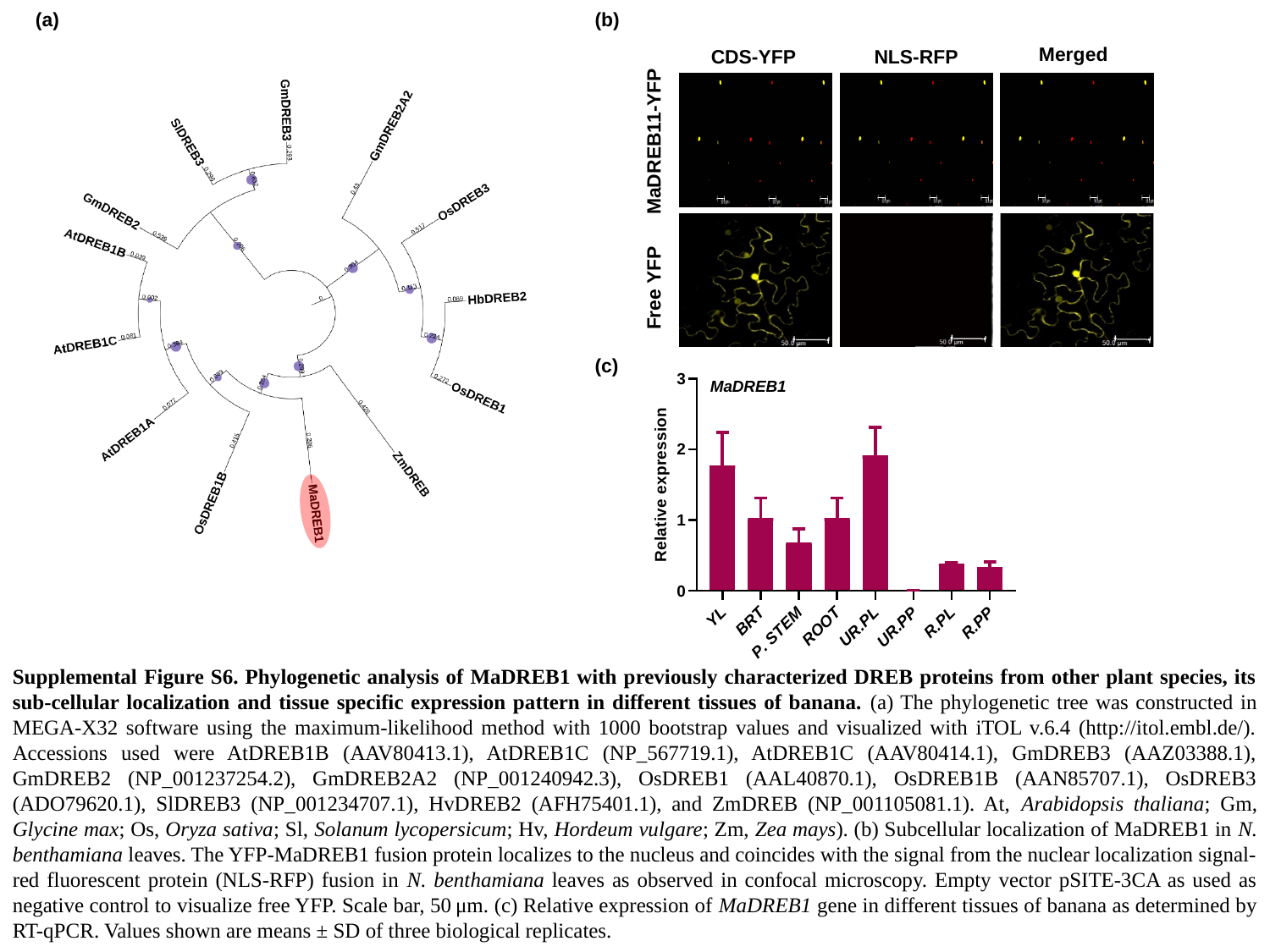

(a)
(b)
Merged
CDS-YFP
NLS-RFP
MaDREB11-YFP
Free YFP
(c)
Supplemental Figure S6. Phylogenetic analysis of MaDREB1 with previously characterized DREB proteins from other plant species, its sub-cellular localization and tissue specific expression pattern in different tissues of banana. (a) The phylogenetic tree was constructed in MEGA-X32 software using the maximum-likelihood method with 1000 bootstrap values and visualized with iTOL v.6.4 (http://itol.embl.de/). Accessions used were AtDREB1B (AAV80413.1), AtDREB1C (NP_567719.1), AtDREB1C (AAV80414.1), GmDREB3 (AAZ03388.1), GmDREB2 (NP_001237254.2), GmDREB2A2 (NP_001240942.3), OsDREB1 (AAL40870.1), OsDREB1B (AAN85707.1), OsDREB3 (ADO79620.1), SlDREB3 (NP_001234707.1), HvDREB2 (AFH75401.1), and ZmDREB (NP_001105081.1). At, Arabidopsis thaliana; Gm, Glycine max; Os, Oryza sativa; Sl, Solanum lycopersicum; Hv, Hordeum vulgare; Zm, Zea mays). (b) Subcellular localization of MaDREB1 in N. benthamiana leaves. The YFP-MaDREB1 fusion protein localizes to the nucleus and coincides with the signal from the nuclear localization signal-red fluorescent protein (NLS-RFP) fusion in N. benthamiana leaves as observed in confocal microscopy. Empty vector pSITE-3CA as used as negative control to visualize free YFP. Scale bar, 50 μm. (c) Relative expression of MaDREB1 gene in different tissues of banana as determined by RT-qPCR. Values shown are means ± SD of three biological replicates.

### Slide 9
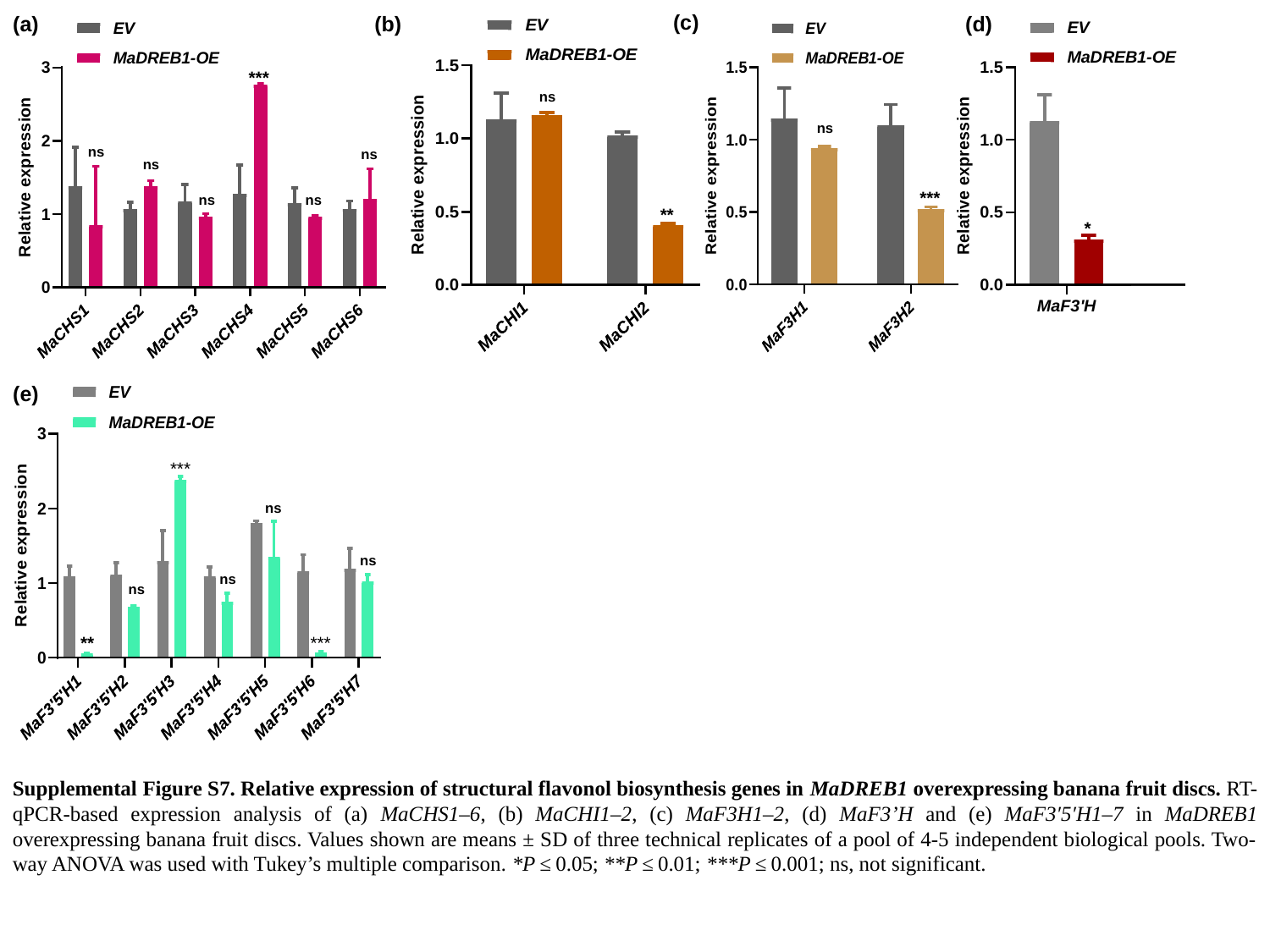

(c)
(b)
(a)
(d)
***
ns
ns
ns
ns
ns
***
ns
ns
**
*
(e)
***
ns
ns
ns
ns
***
**
Supplemental Figure S7. Relative expression of structural flavonol biosynthesis genes in MaDREB1 overexpressing banana fruit discs. RT-qPCR-based expression analysis of (a) MaCHS1–6, (b) MaCHI1–2, (c) MaF3H1–2, (d) MaF3’H and (e) MaF3′5′H1–7 in MaDREB1 overexpressing banana fruit discs. Values shown are means ± SD of three technical replicates of a pool of 4-5 independent biological pools. Two-way ANOVA was used with Tukey’s multiple comparison. *P ≤ 0.05; **P ≤ 0.01; ***P ≤ 0.001; ns, not significant.

### Slide 10
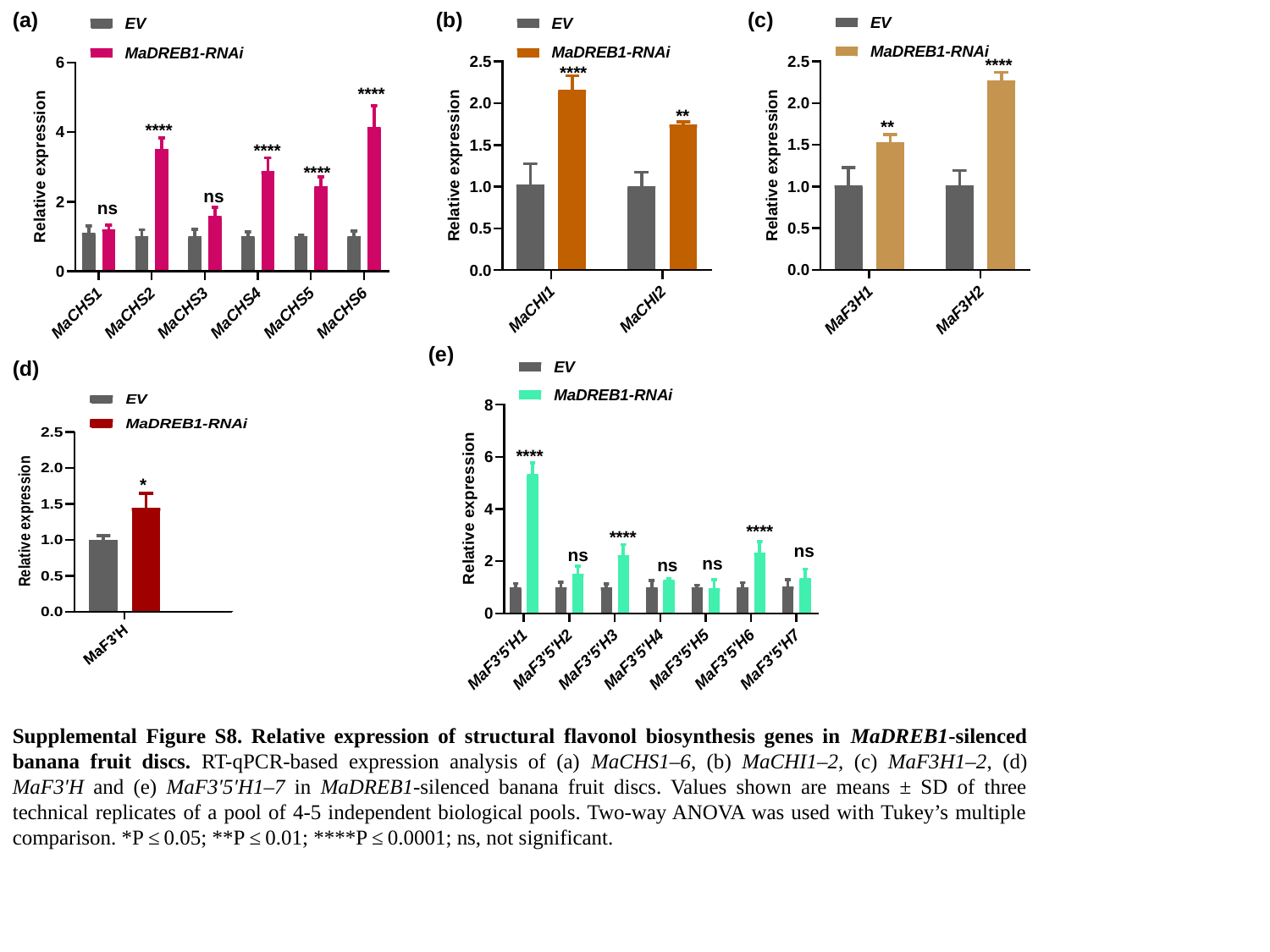

(a)
(b)
(c)
****
****
****
**
**
****
****
****
ns
ns
(e)
(d)
****
*
****
****
ns
ns
ns
ns
Supplemental Figure S8. Relative expression of structural flavonol biosynthesis genes in MaDREB1-silenced banana fruit discs. RT-qPCR-based expression analysis of (a) MaCHS1–6, (b) MaCHI1–2, (c) MaF3H1–2, (d) MaF3′H and (e) MaF3′5′H1–7 in MaDREB1-silenced banana fruit discs. Values shown are means ± SD of three technical replicates of a pool of 4-5 independent biological pools. Two-way ANOVA was used with Tukey’s multiple comparison. *P ≤ 0.05; **P ≤ 0.01; ****P ≤ 0.0001; ns, not significant.

### Slide 11
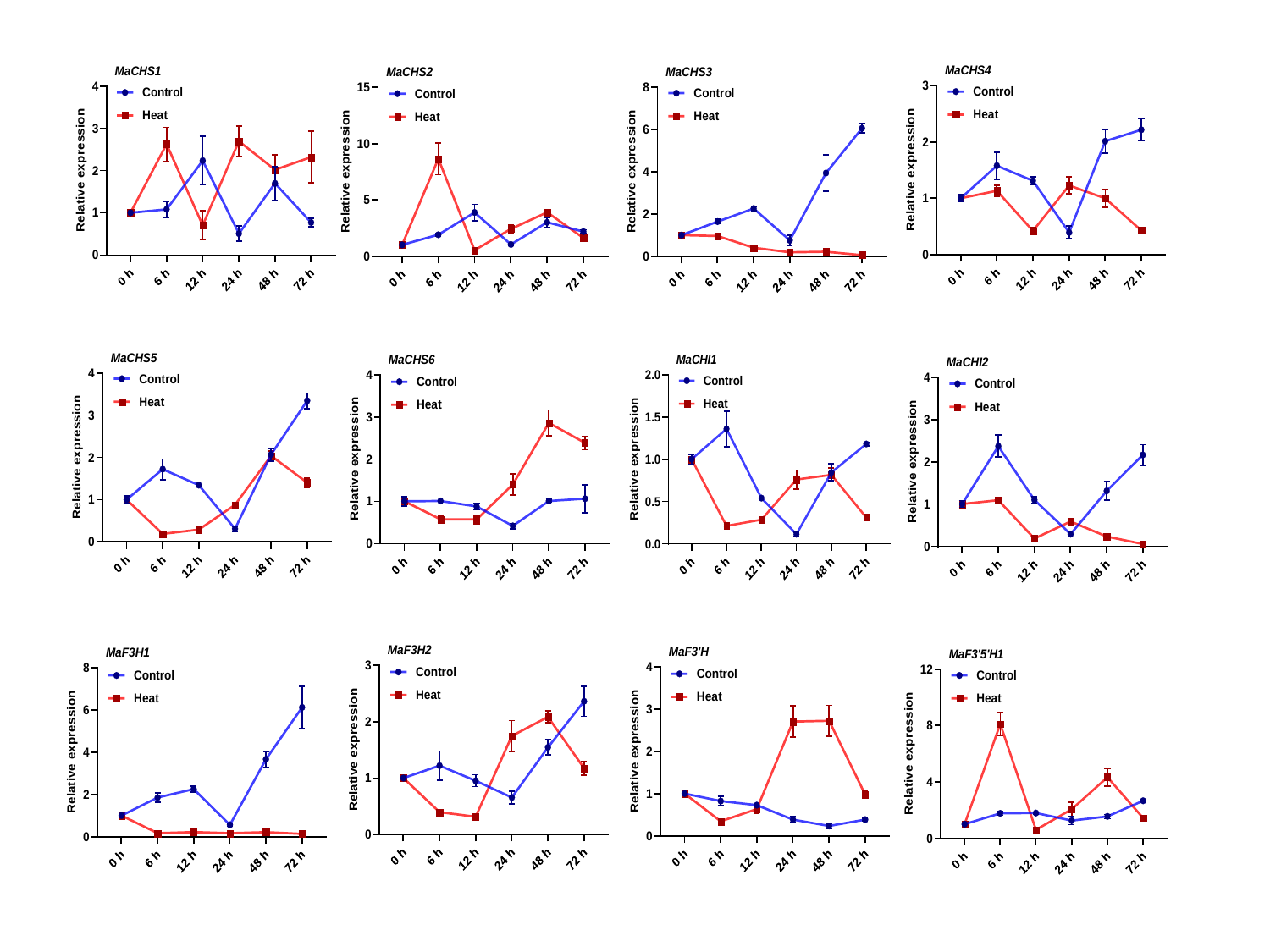

### Slide 12
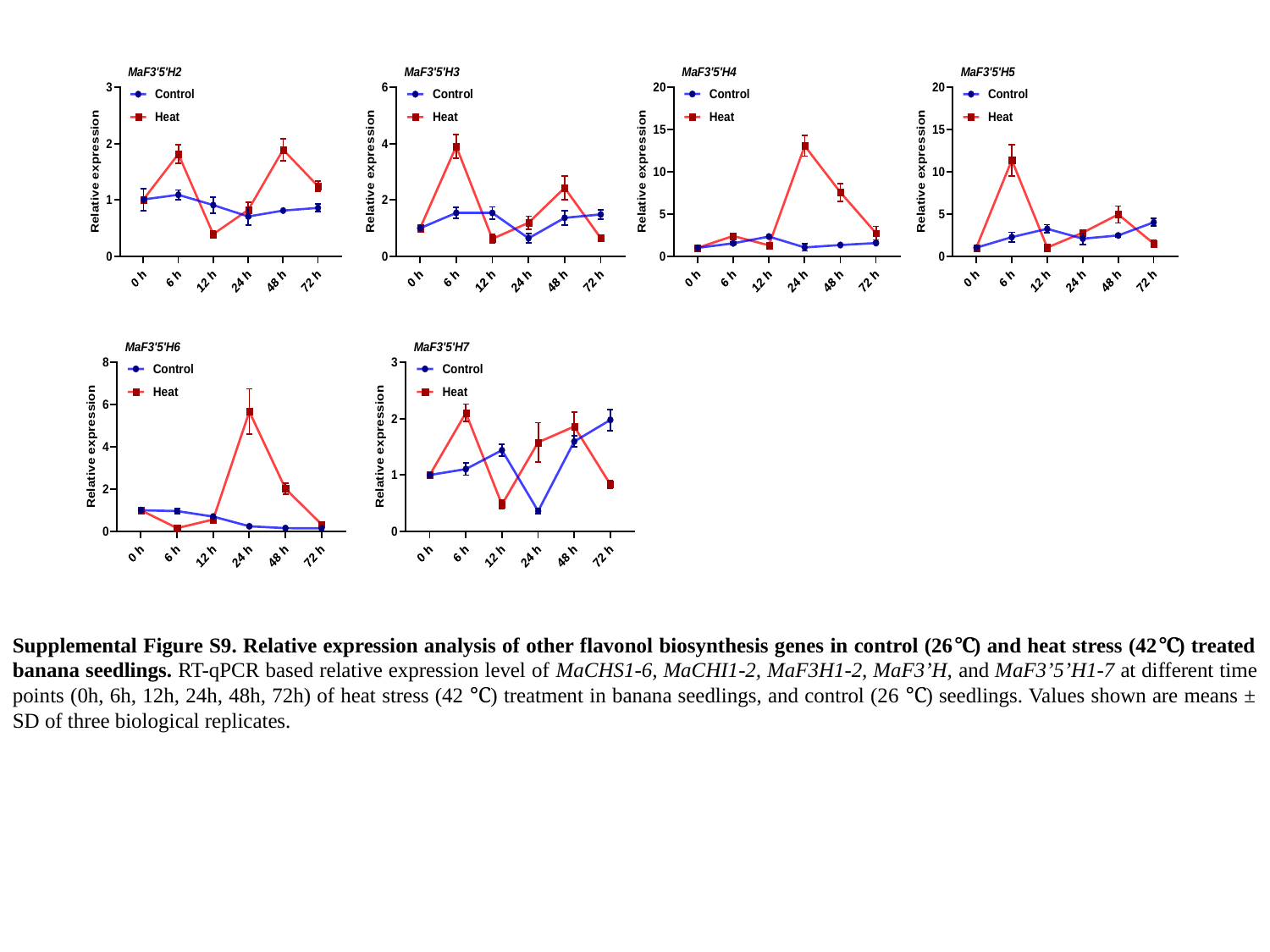

Supplemental Figure S9. Relative expression analysis of other flavonol biosynthesis genes in control (26℃) and heat stress (42℃) treated banana seedlings. RT-qPCR based relative expression level of MaCHS1-6, MaCHI1-2, MaF3H1-2, MaF3’H, and MaF3’5’H1-7 at different time points (0h, 6h, 12h, 24h, 48h, 72h) of heat stress (42 ℃) treatment in banana seedlings, and control (26 ℃) seedlings. Values shown are means ± SD of three biological replicates.

### Slide 13
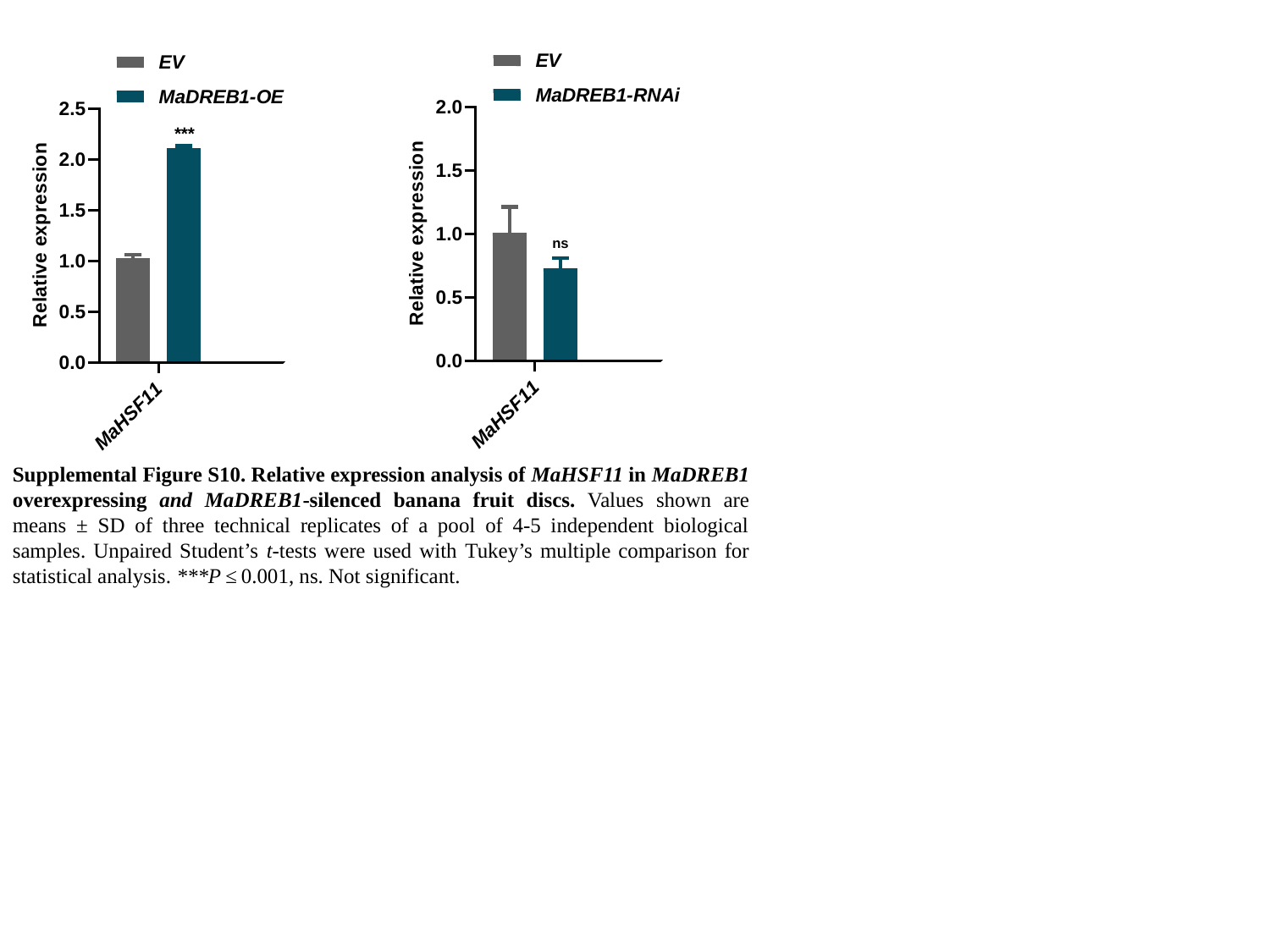

***
ns
Supplemental Figure S10. Relative expression analysis of MaHSF11 in MaDREB1 overexpressing and MaDREB1-silenced banana fruit discs. Values shown are means ± SD of three technical replicates of a pool of 4-5 independent biological samples. Unpaired Student’s t-tests were used with Tukey’s multiple comparison for statistical analysis. ***P ≤ 0.001, ns. Not significant.

### Slide 14
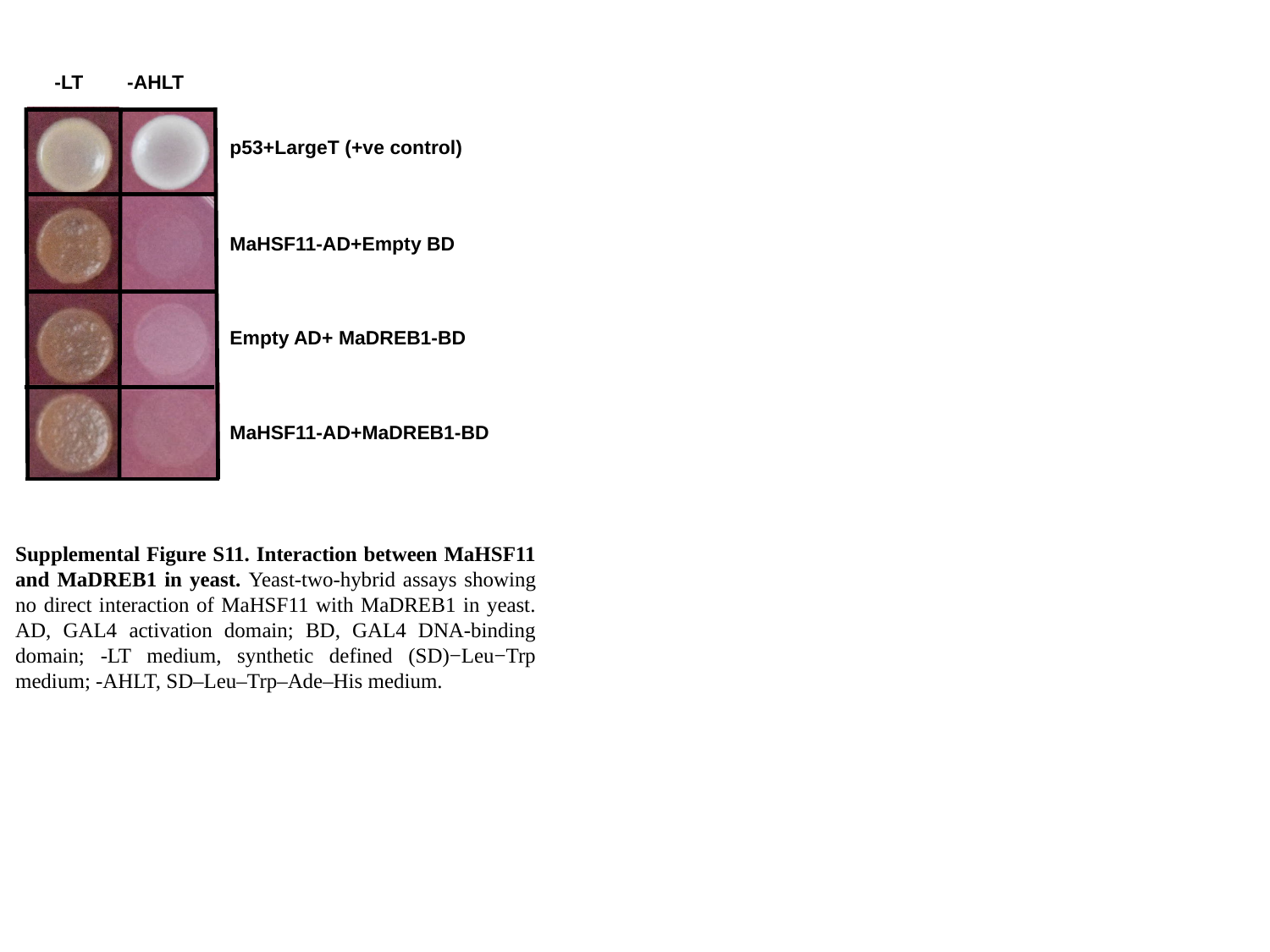

-LT -AHLT
p53+LargeT (+ve control)
MaHSF11-AD+Empty BD
Empty AD+ MaDREB1-BD
MaHSF11-AD+MaDREB1-BD
Supplemental Figure S11. Interaction between MaHSF11 and MaDREB1 in yeast. Yeast-two-hybrid assays showing no direct interaction of MaHSF11 with MaDREB1 in yeast. AD, GAL4 activation domain; BD, GAL4 DNA-binding domain; -LT medium, synthetic defined (SD)−Leu−Trp medium; -AHLT, SD–Leu–Trp–Ade–His medium.
